# Spatiotemporal Trajectories in Resting-state FMRI Revealed by Convolutional Variational Autoencoder

**DOI:** 10.1101/2021.01.25.427841

**Authors:** Xiaodi Zhang, Eric Maltbie, Shella Keilholz

## Abstract

Recent resting-state fMRI studies have shown that brain activity exhibits temporal variations in functional connectivity by using various approaches including sliding window correlation, co-activation patterns, independent component analysis, quasi-periodic patterns, and hidden Markov models. These methods often model the brain activity as a discretized hopping among several brain states that are defined by the spatial configurations of network activity. However, the discretized states are merely a simplification of what is likely to be a continuous process, where each network evolves over time following its unique path. To model these characteristic spatiotemporal trajectories, we trained a variational autoencoder using rs-fMRI data and evaluated the spatiotemporal features of the latent variables obtained from the trained networks. Our results suggest that there are a relatively small number of approximately orthogonal whole-brain spatiotemporal patterns that capture the most prominent features of rs-fMRI data, which can serve as the building blocks to construct all possible spatiotemporal dynamics in resting state fMRI. These spatiotemporal patterns provide insight into how activity flows across the brain in concordance with known network structures and functional connectivity gradients.

## 1 Introductions

In resting state fMRI (rs-fMRI), the blood oxygenation level-dependent (BOLD) signal is acquired in the absence of an explicit task or stimulation (Biswal et al., 1995; Ogawa et al., 1992). Networks of spatially distributed brain regions whose time courses are correlated, referred to as “resting state networks” (RSN) (Cordes et al., 2000; Damoiseaux et al., 2006; Fox et al., 2006; Fox and Raichle, 2007; Ghahremani et al., 2016; Greicius et al., 2003; Hampson et al., 2002; Power et al., 2011; Smith et al., 2009), can be reliably observed under numerous conditions and serve as the foundation of our knowledge of the brain’s functional architecture. Recent studies have revealed that these large-scale patterns of brain activity exhibit temporal variations at relatively fast time-scales (seconds-minutes) (Allen et al., 2014; Chang and Glover, 2010; Handwerker et al., 2012; Jones et al., 2012a; Keilholz et al., 2013; Kiviniemi et al., 2011; Majeed et al., 2011; Sakoğlu et al., 2010), and that these dynamics are sensitive to changes related to behavior, cognition (Albert et al., 2009; Bassett et al., 2011; Esposito et al., 2006; Fornito et al., 2012; Thompson et al., 2013), and pathology (Damaraju et al., 2014; Hamilton et al., 2011; Jones et al., 2012a). A number of techniques have been used to characterize the time-varying patterns of activity, including sliding window correlation (SWC) (Allen et al., 2014; Chang and Glover, 2010; Handwerker et al., 2012; Jones et al., 2012a; Keilholz et al., 2013; Kiviniemi et al., 2011), co-activation patterns (CAPs) (Liu and Duyn, 2013; Tagliazucchi et al., 2012), Independent component analysis (ICA) (Allen et al., 2014; Damaraju et al., 2014; Kiviniemi et al., 2011) and hidden Markov models (HMM) (Vidaurre et al., 2017). However, most of these methods consider spatial and temporal information separately, when in reality the temporal and spatial aspects of brain activity are intricately related. Brain activity has often been modeled as a discretized hopping among several brain states that are defined by the spatial configurations of network activity. However, the discretized states are merely a simplification of what is likely to be a continuous process, where each network evolves over time following its unique path. In this case, the presence of stereotyped pathways of evolution between states that manifest as characteristic spatiotemporal trajectories in the rs-fMRI data would provide new insight into the systems-level coordination of brain function.

At least one characteristic spatiotemporal trajectory has already been observed using a recursive algorithm. The resulting quasi-periodic patterns (QPPs) revealed highly reproducible spatiotemporal trajectories showing sinusoidal patterns of activation and deactivation in the default mode network (DMN) and task positive network (TPN) with opposite phases (Abbas et al., 2019; Yousefi et al., 2018), along with propagation along the cortex. Despite these successes, the primary QPP only explains 25-50% of the variance in the BOLD signal (Hutchison et al., 2013), suggesting that there is still a large portion of the signal not accounted for, and there are potentially other spatiotemporal trajectories not yet identified. An effort has been made to identify these secondary components by performing QPP analysis again after regressing out the primary QPP component (Belloy et al., 2018). These secondary QPPs have demonstrated distinct spatiotemporal patterns that are different from the primary ones, however the number of additional components identified was limited to three. To date there has not been an exhaustive search for all possible characteristic spatiotemporal trajectories, potentially due to difficulties from the computational complexities, as well as the reduced robustness and interpretability after the repeated calculation of regression and convolution.

Deep learning methods could potentially solve this problem because they are inherently designed to extract key information or characteristic patterns from very complicated systems in a data-driven way. Convolutional neural networks (CNN), in particular,) have proven very successful at extracting spatial features from images, e.g. AlexNet (Krizhevsky et al., 2017) and GoogLeNet (Szegedy et al., 2015), and there are also studies using convolutional neural networks to extract temporal features from time series, e.g. applications in natural language processing (Gehring et al., 2017; Kalchbrenner et al., 2014; Kim, 2014) where the convolutional kernel was shown to be capable of extracting the features from the ordering of words in a sentence. In a more generic setting, (Bai et al., 2018) has shown that the CNN is capable of learning the temporal structures of time series in various tasks. Therefore, supposing there is a specific spatiotemporal property attributable to intrinsic brain dynamics, presumably it would be captured by a CNN as well.

As of today there are relative few studies in resting state fMRI that use deep learning methods, most of which focus on classification problems, e.g., classification of Alzheimer’s disease (Sarraf and Tofighi, 2017), mild cognitive impairment (MCI) (Meszlényi et al., 2017; Suk et al., 2016) and ADHD (Mao et al., 2019). A few studies attempt to extract features in the fMRI data. For example, Huang et al. (2018) used a convolutional autoencoder to extract temporal features from task-fMRI data, which describes variations in the hemodynamic response function (HRF). Hu et al. (2018) trained a restricted Boltzmann machine using task-fMRI data, which was claimed to outperform ICA in terms of higher temporal correlation with task paradigms, and greater spatial overlap with the general linear model. Despite deep learning’s great potential, none of the existing studies is designed to detect characteristic spatiotemporal brain trajectories.

We designed a deep learning method specifically to extract characteristic spatiotemporal trajectories from rs-fMRI time courses. Specifically, a variational autoencoder (VAE) was trained to identify a relatively small number of approximately orthogonal whole-brain spatiotemporal patterns that capture the most prominent features of rs-fMRI data. The resulting latent variables show that characteristic brain trajectories (beyond the QPP) exist and provide insight into how activity flows across the brain in concordance with known network structures and functional connectivity gradients.

## 2 Methods

### 2.1 fMRI data preprocessing

The minimally processed rs-fMRI data from the 412 subjects with “study completion: full 3T imaging protocol completed” label was downloaded from the HCP S500 release (Glasser et al., 2013). The resting-state fMRI data were acquired using Gradient-echo EPI with the following parameters: TR/TE = 720ms/33.1ms, resolution = 2.0mm isotropic, matrix size = 104×90, number of slice = 72, numer of TR = 1200. Further preprocessing included the following procedures: The first 5 frames were removed to minimize the transient effects before reaching equilibrium. Gray matter (GM), white matter (WM) and cerebrospinal fluid (CSF) signal were averaged within their masks provided by HCP. Then GM, WM, and CSF signals, along with 12 motion parameters (provided by HCP), linear and quadratic trends were regressed out altogether at the voxel level. The regressed BOLD signals were then bandpass filtered using a 0.01-0.1Hz 6-order Butterworth filter, and spatially smoothed using a Gaussian kernel with FWHM = 4mm. Finally the BOLD signals were parcellated using the Brainnetome atlas (Fan et al., 2016) and each parcel was z-scored. The final parcellated BOLD signal has 412 subjects by 1195 time points by 246 parcels. For better visualization, the 246 parcels were then sorted into 7 functional networks using Yeo’s 7-network model (Thomas Yeo et al., 2011) provided by the Brainnetome website, namely default mode (DMN), visual (VIS), somatomotor (SM), dorsal attention (DA), ventral attention (VA), frontalparietal (FP) and limbic (LIM) networks, with the remaining parcels all classified as subcortical regions (SC).

### 2.2 Variational Autoencoder

An autoencoder is a type of neural network used to learn efficient data representation in an unsupervised manner. It typically consists of an encoder network that gradually reduces dimensions, and a symmetric decoder network that recovers the dimensions. In this case, the output of the encoder has the lowest dimensionality in the entire network, and thus is a bottleneck of the information, which forces the network to extract features that most represent the data structure, since any reconstruction error is penalized.

To improve generalizability, a variant of the autoencoder architecture called a variational autoencoder (VAE) includes a random sampling process (Kingma and Welling, 2014). The model learns the distributions of the latent variables (by learning means and variances), instead of learning a deterministic mapping. A random sample is drawn from the distributions for every data point passing through the latent layer. The calculation of the loss function involved in this process and how it is back propagated to update the parameters in the networks was described in the original VAE paper (Kingma and Welling, 2014). To summarize, the loss function that corresponds to the randomization process is the Kullback-Leibler (KL) divergence, which has a closed form when the prior distribution is assumed to be Gaussian. Thus, by minimizing the sum of the reconstruction loss and the KL divergence, the latent variable not only learns the most representative features in the dataset, but also becomes as close to a multidimensional standard Gaussian distribution (all components are independent, zero-mean, unit-variance) as possible. This tendency to approach Gaussian distribution serves as a regularization effect, which leads to a smoother latent distribution compared to the plain autoencoder, and thus improves the generalizability of the model. The VAE model essentially assumes that if the network is deep enough (having enough expressive power), then any complicated system can be mapped to a series of disentangled Gaussian-distributed variables.

### 2.3 Convolutional Variational Autoencoder Design

With the goal of extracting common spatiotemporal trajectories in brain activities, we chose to feed the neural network with short rs-fMRI segments instead of single frames. Each rs-fMRI scan (1195 TR) was divided into 36 segments that are 33-TR long (23.76sec), with 50% overlap. The 33-TR segment length was chosen based on prior work identifying a strong spatiotemporal pattern with a duration of ~20s (Majeed et al., 2011). Based on the assumption that the rules governing the network dynamics are shift-invariant across time, convolutional layers were used in the first few layers instead of fully connected layers. As suggested by (Lecun et al., 1998), the parameter sharing in the convolutional layer greatly reduces the number of parameters in the model, thus improves its generalizability. Instead of using the common 2D convolutional kernel, here we used a 1D convolutional kernel that applies only to the temporal dimension, because the fMRI signal in the parcellated space is not shift-invariant across different parcels in the spatial domain.

This neural network architecture is shown in figure 1. The network consists of a symmetric encoder and decoder pair, either of which has 3 convolutional layers and 2 fully-connected layers. Each convolutional/fully-connected layer consists of a weight layer and a Rectified Linear Unit (ReLU) activation layer. The performance of other 4 alternative network designs with different number of layers or different number of hidden units was evaluated using holdout validation (the results are shown in supplemental materials section S.1) and the architecture shown in figure 1 showed the best performance. The encoder encodes the input rs-fMRI segments of size 246 parcels by 33 time points into a 32×1 latent representation that roughly follows a multidimensional Gaussian distribution. The distributions of the latent variables were represented in means and variances that are estimated by the networks. Then during training, a sample was randomly drawn from this distribution whenever a data point arrives at the latent layer. This random process is a key feature in variational autoencoder, which improves its robustness and generalizability. Then the decoder performs a series of reverse operations (dilated convolution being the reverse operation of convolution) to reconstruct rs-fMRI segments from the 32×1 latent representation.

**Figure 1.**
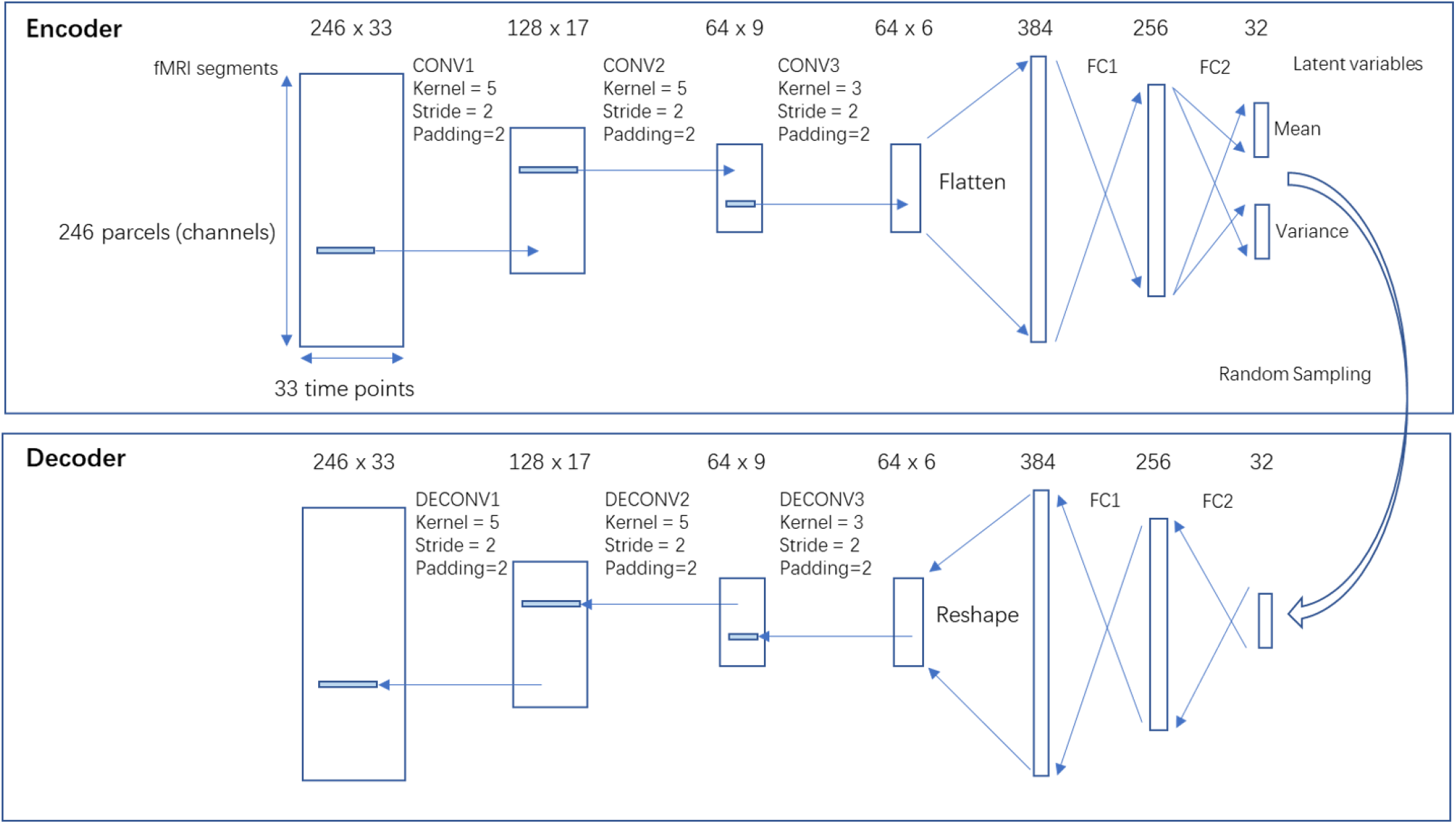
The architecture of the networks. The networks consist of a symmetric encoder and decoder, both having 3 convolutional or dilated convolutional layers, and 2 fully connected layers. The encoder encodes rs-fMRI segments of size 246×33 into 32×1 latent variables that follow Gaussian distributions, whose mean and variance were estimated by the network. Then a sample is randomly drawn from the distribution, which is then propagated through the decoder to reconstruct back to rs-fMRI segments.

### 2.4 Training and testing of the model

The 412 subjects were randomly split into a training set (n=248), a validation set (n=82) and a testing set (n=82). Then the segments were shuffled, resulting a training set with size of [248×36,246,33], a validation set and a testing set both with size of [82×36,246,33]. To make the model more regularized, we used a variant of VAE called beta-VAE (Higgins et al., 2017), whose loss function is the sum of reconstruction loss (root mean square error between input and output) and the K-L divergence loss weighted by a factor beta (beta=4). Large beta values increase the penalty for KL-divergence and therefore the model is more regularized (variables become closer to orthogonal). As proposed in the original beta-VAE paper, as well as confirmed in our experiments (shown in supplemental materials section S.2), beta = 4 gives a reasonable result that appear to be more robust and regularized than a regular VAE (a special case where beta = 1). The networks were trained on a Nvidia GTX2080Ti GPU using Adam optimizer (Kingma and Ba, 2015) with a learning rate of 0.001 for 90 epochs. To validate the model, we used the rs-fMRI segments from the testing set as the input and compared the rs-fMRI segments reconstructed by the networks with the input. The reconstruction provides a qualitative assessment of how much information is preserved by the latent representation.

### 2.5 Feature Visualization of the Latent Variables

Neural networks are often described as “black boxes” and it is not uncommon to see difficulties in interpreting why they perform well over a particular task. There are a few methods for visualizing features learned by the networks that can help interpret the results, including saliency maps and class visualization (Simonyan et al., 2014), although these methods are typically used for classifiers. Thanks to its Gaussian-distributed latent variables and its symmetric encoder-decoder design, there is one visualization method exclusive to variational autoencoder. The latent variables are disentangled, because penalizing the KL divergence leads to a multidimensional Gaussian distribution where all components are independent from each other. This means that the effect of each latent variable is isolated, thus can be visualized by propagating a perturbation of such latent variable though the decoder. In addition, this effect on the reconstructed rs-fMRI segments by the decoder, should ideally be the same spatial temporal pattern that will activate the corresponding latent variable when passing through the encoder. Using this method, we vary each of the 32 latent variables from −3 to +3 (since 99.7% of the data lies in the ±3 sigma range of a Gaussian distribution) with 500 increment steps, and observe how the rs-rs-fMRI segments reconstructed by the decoder vary. This process returns a 4D vector (500 increments, 246 parcels, 33 time points, 32 latent variables), which can be visualized if one dimension is fixed. By fixing the perturbation at its maximum amplitude, we obtained a set of 32 spatiotemporal patterns or trajectories of brain activities that can activate their corresponding latent variables, which is shown in figure 2.

**Figure 2.**
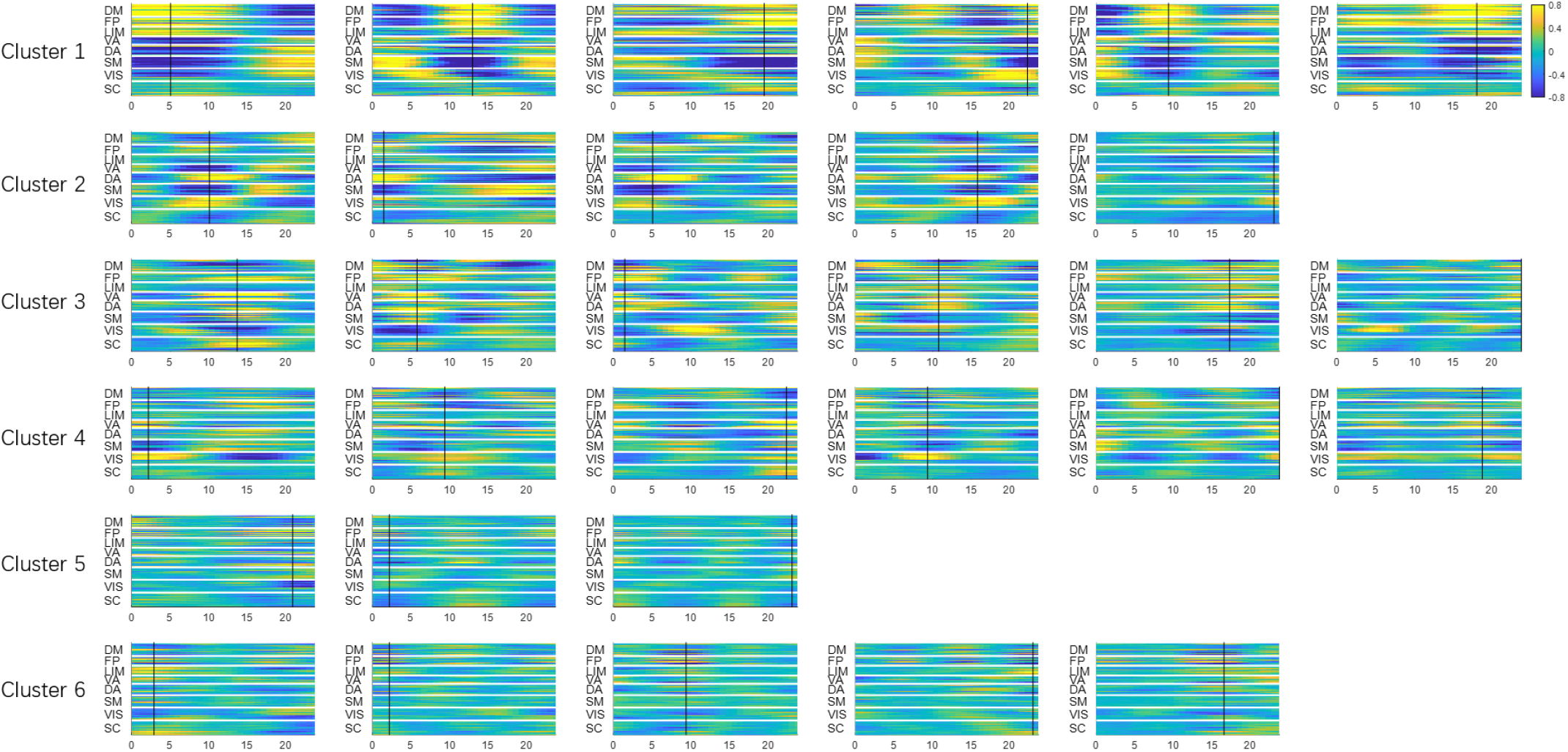
Spatiotemporal patterns extracted by latent variables. Each subplot is obtained by making one latent variable equal to +3 (corresponding to +3σ for a Gaussian distribution) while fixing the rest of the latent variables at zero. The x-axis is time in seconds. The y-axis is the 246 parcels. The patterns have arbitrary units, but all subplots share the same display scale so that higher variance results in higher contrast. The 32 latent variables are already organized in 6 clusters (see their spatial configurations in figure 3). The black cursor indicates the time of maximum spatial variance across parcels.

### 2.6 Grouping of the Latent Variables Based on Their Spatial Similarity

These 32 spatiotemporal patterns exhibit a few common spatial configurations which show synchronized fluctuations. Thus the 32 latent dimensions can be further organized into several groups based on their similarity in the spatial domain. To do that, first the time points when the fMRI time course reaches maximum variance across spatial dimensions were extracted (shown with black cursors in figure 2). The spatial profiles (as a function of latent variable) at the max-variance time points of the 32 latent variables were compared with each other and reorganized into several groups using K-means clustering (with spatial similarity calculated with Pearson correlation being the clustering criteria, and k empirically chosen as 6).

Then clusters were sorted in descending order by the variance explained by each latent variable (calculated as the variance across time domain, which was then summed over 246 parcels). The variance of individual latent variables within a cluster is also in descending order for better visualization. Aside from the spatial profiles, the functional connectivity of each latent variable’s spatiotemporal pattern was calculated. The weighted average (weighted by the variance of the latent variable) functional connectivity within each cluster was shown to provide an alternative representation of the spatial configurations among major functional networks of the 6 clusters.

### 2.7 Comparison with the Primary QPP

The latent variables of the trained networks capture spatiotemporal trajectories of the brain, in a manner similar to the QPPs. Thus the features of latent variable 1, whose variance is the highest, was compared with the primary QPP. The primary QPP was calculated from the same testing set (n=82) with the Brainnetome parcellation, using the existing Matlab code for calculating QPPs published in (Yousefi et al., 2018).

## 3 Results

### 3.1 The convolutional VAE decomposes rs-fMRI segments into a weighted combination of spatiotemporal patterns

The trained convolutional VAE learns to represent any rs-fMRI segments using the 32 latent variables. To visualize the latent variables, we used the method described in section 2.5. Figure 2 shows a set of 32 spatiotemporal trajectories of brain activity that can activate their corresponding latent variables. This set of spatiotemporal patterns were learnt to be the most representative features existing in short rs-fMRI segments, and any given rs-fMRI segment can be expressed by a weighted sum of these orthogonal spatiotemporal patterns, with the weights being the values of latent variables for that particular rs-fMRI segment. Note that each cluster of the spatiotemporal trajectories shares a common spatial network configuration (which can also be seen in the clusters in figure 3), while each individual latent variable within a given cluster describes a unique evolution of activity for that particular network configuration. These latent variables are organized into 6 groups based on their spatial configurations using the method described in section 2.6. It can be seen that each cluster shares a common spatial organization of connectivity. For example, all 6 of the latent variables in the first cluster exhibit the anticorrelated DMN-TPN network configuration. All 32 spatiotemporal patterns share the same display scale, thus higher contrast suggests higher variance explained and presumably greater importance of the latent variable.

**Figure 3.**
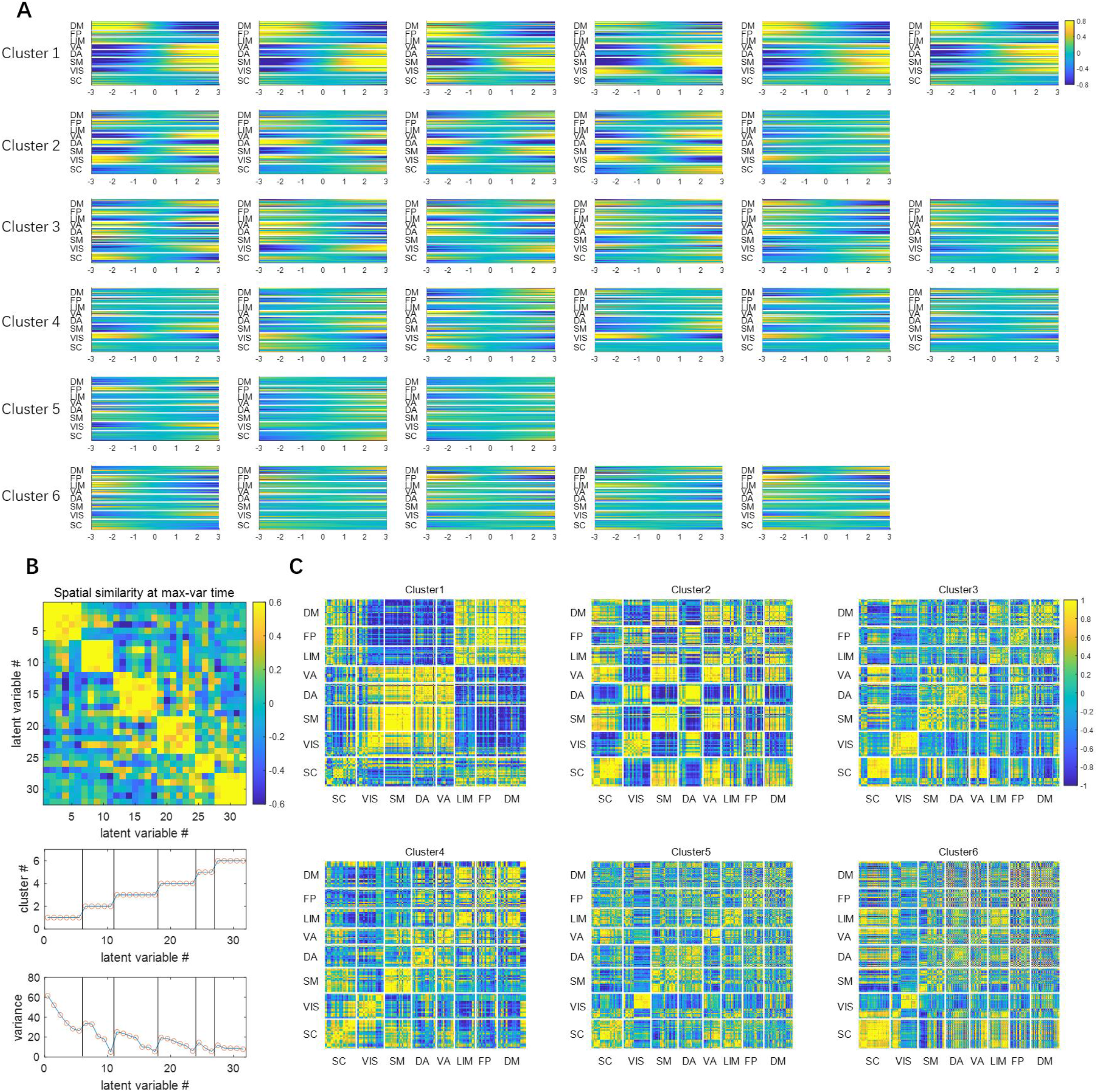
The latent dimensions can be organized into 6 clusters (shown in rows) based on their spatial similarities. Panel A shows how the spatial profile at the max-variance time (in figure 2) changes when sliding a single latent variable from −3 to +3. Panel B shows the spatial similarities among latent variables at the max-variance time, measured by Pearson correlation between the spatial profiles. The latent variables were then clustered using K-means clustering using the spatial similarity as the clustering criteria (K = 6). The cluster label index and the variance explained are also shown. Panel C shows the weighted mean functional connectivity of each cluster of latent variables over the 33-TR window.

### 3.2 The 32 latent dimensions can be further clustered based on their spatial similarity

To better illustrate the common spatial configurations shared by the latent variables, here we leave out the temporal dimension by focusing on the time point when the fMRI time course reaches maximum variance across spatial dimensions, as described in section 2.6. The spatial configurations at this timepoint are shown for each variable in each cluster in Figure 2, accompanied by a matrix of the spatial similarity (Pearson correlation) between the spatial configurations that clearly shows the division into six distinct groups. The weighted averaged functional connectivity for each group is also shown to provide an alternative representation of the spatial configurations, and the variance explained for each latent variable is given.

It can be seen in figure 3 panel A that, within the primary cluster, whose mean variance is the highest, the spatial profile of every latent dimension at the max-variance time has the DM, FP and LIM network on one end, and VIS, SM, DA and VA networks on the opposite end. Although this max-variance time only gives a snapshot of this opposing relationship, such contrast can be seen throughout the course of the trajectories (both shown in the time courses in figure 2, and the functional connectivity in figure 3 panel C). This finding is in agreement with many previous studies, including the DMN/TPN anticorrelation found in (Fox et al., 2005), quasiperiodic patterns (Majeed et al., 2011) and principal functional connectivity gradients (Margulies et al., 2016). The latent variables in the primary cluster all show that the DMN and TPN have a few components (with very high variance) with opposite phase at almost every instantaneous moment, suggesting this is the most prominent feature existing in resting state fMRI, which is likely the reason why we can see a consistent anti-correlation between the two networks.

The secondary cluster, which has the second highest variance, also has an interesting feature that further separates different networks within the task positive network. At the max-variance time, it can be seen from figure 3 panel A that, every latent variable in cluster 2 has the negative end corresponding to the activation of VIS and DA networks, and the positive end corresponding to the activation of SM and VA networks. These together with the primary cluster, exhibit a remarkable resemblance to the principal gradients. The principal gradients are obtained using a method called diffusion embedding, which maps brain regions into an embedded space, where strongly connected points are closely spaced while loosely connected points are far apart. It was reported that in principle gradient 1, the transmodal DMN regions are anchored at one end and the unimodal visual, somatosensory/motor regions are at the other end, whereas in principle gradient 2, the visual networks are at one end and the somatosensory/motor regions are on the opposite end. This close resemblance between latent variables and principal gradients provides evidence that the network configurations based on the connectivity geometry revealed by the principal gradients closely reflects the instantaneous network activity demonstrated by the VAE.

### 3.3 The primary latent dimension shows a spatial-temporal pattern very similar to the QPP

It can be seen from figure 2 that the first latent dimension (which has the highest variance) encodes a spatiotemporal pattern that shows one cycle of anti-correlated activities between DMN and TPN over a 24 second time window. This spatiotemporal feature is very similar to the primary QPP except having opposite phase (the phase in figure 4 is already reversed for better comparison with QPP). The network is trained with randomly initialized weights, which leads to random polarity of latent variables for every training trial. Thus, the polarity can be ignored and the latent variable 1 and the primary QPP essentially extracted very similar information. This similarity makes sense because QPP averages the time points that have the most prominent correlation with the template, thus reinforcing itself over multiple iterations, and extracting the most prominent, reoccurring spatial temporal features. It is not surprising that such spatial temporal features have the most variance and thus were picked up by the variational autoencoder as the first latent dimension.

**Figure 4.**
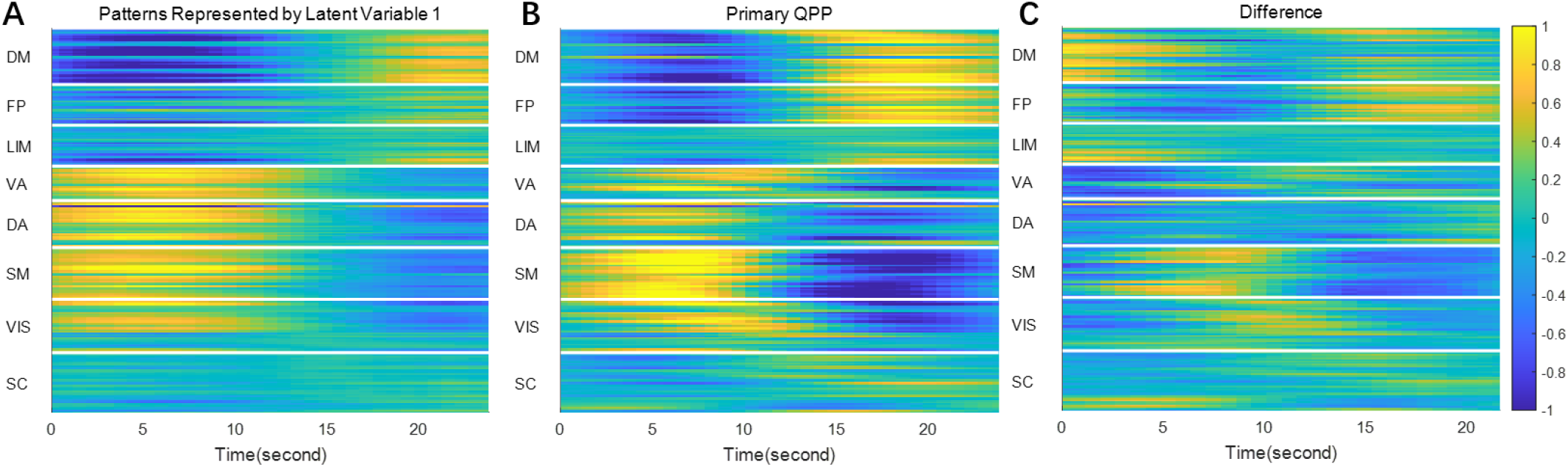
Spatial temporal features represented by latent variable 1 (panel A), the primary QPP (panel B) and their difference. Both the latent feature and the QPP were divided by their 98th percentile to normalize. It can be seen that the spatial temporal features represented by latent variable 1 are very similar to the primary QPP (Pearson correlation coefficient = 0.759), but there are also some differences, most notably in the strength of frontoparietal involvement and near transitions between positive and negative activation in the somatomotor network.

Aside from the first latent dimension, there are also 5 other latent dimensions in the primary cluster that share very similar spatial distributions, but differ in frequency and phase. To better visualize these differences among the timings of the latent variables, the latent features from a region of interest (ROI) in the SM was shown as a function of both the value of latent variable and time in figure 5. Specifically, these latent variables with smaller variance tend to have higher frequencies. These spatiotemporal trajectories have not been previously reported, probably because their variance is relatively small compared to the primary component. On top of this, the VAE also identifies 5 other clusters of latent variables that have different spatial configurations. In the traditional QPP calculations, these features may have been canceled out with each other during the averaging process.

**Figure 5.**
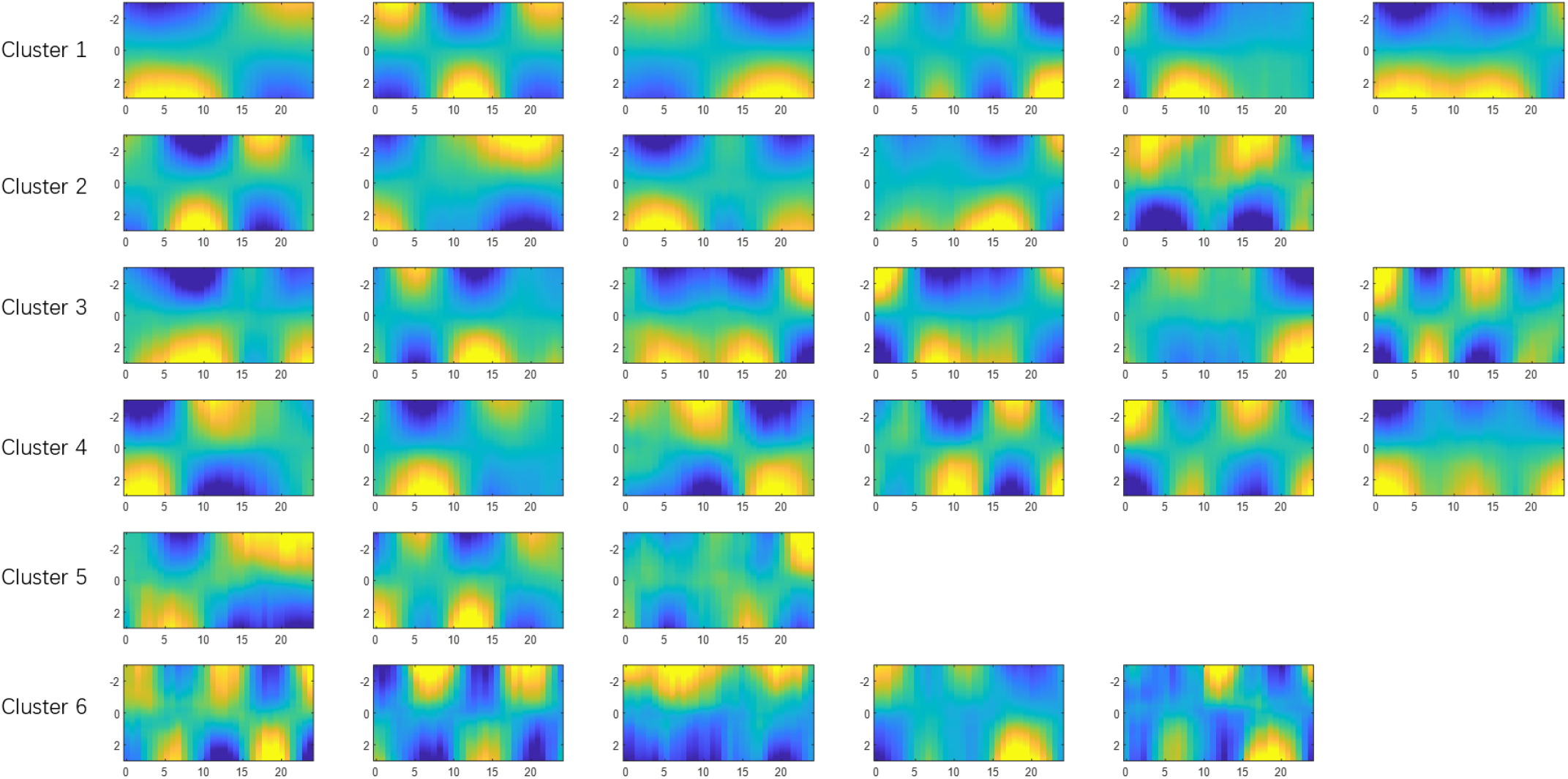
Temporal patterns extracted by latent dimensions. Each subplot shows the temporal pattern from a latent variable obtained by averaging 5 parcels in the SM network (96th parcel to 100th parcel, all in the Postcentral Gyrus). The x-axis is time in seconds. The y-axis is the value of the latent variable, sliding from −3 to +3.

### 3.4 Reconstruction of rs-fMRI segments in the testing set

Figure 6 shows the reconstruction of the rs-fMRI segments and the corresponding weights of latent variables. This reconstruction provides a qualitative assessment of how much information is lost during the encoding-decoding process. Although it is not a perfect match, most of the timing and the amplitude information is captured, especially for fluctuations with high amplitudes. It is worth mentioning that each rs-fMRI segment has 246 parcels and 33 time points, while the encoded representation only has 32 variables, which is around 1/250 of the original size. This fairly good quality of reconstruction despite such a high compression rate suggests that the original parcellated rs-fMRI data is actually quite redundant, which is potentially due to the fact that many parcels coactivate with each other, while others may show anticorrelations. This interlinked relationship among different brain regions greatly reduces the degree of freedom in the system. Thus, the proposed VAE extracts a set of orthogonal bases that accounts for most of the degree of freedom (that have the highest variances), which creates a parsimonious representation of brain activity that reveals such relationships among brain regions.

**Figure 6.**
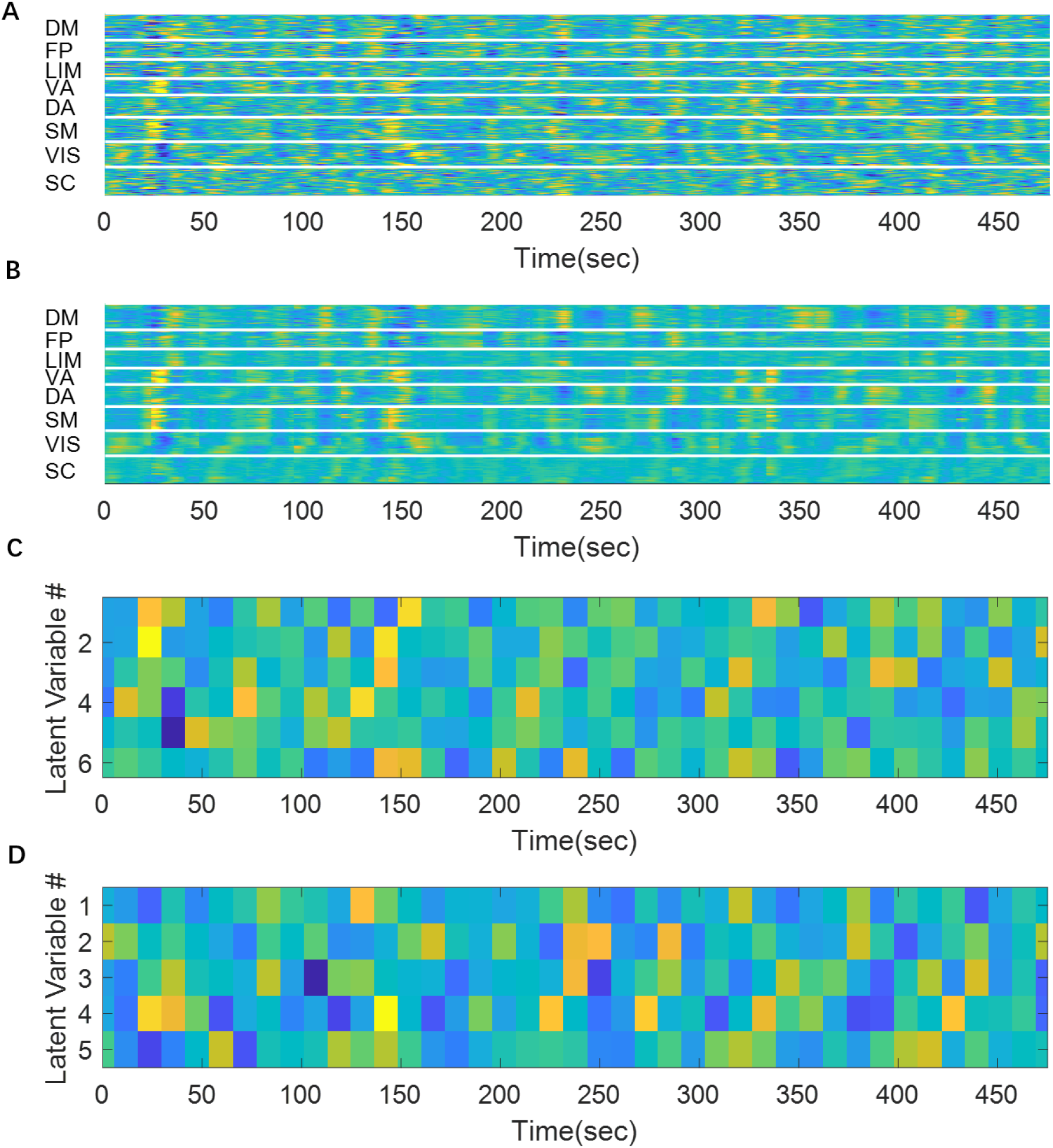
A fMRI segment can be encoded as a 32-dimensional code. Panel A shows 20 concatenated original rs-fMRI segments. Panel B shows the reconstructed rs-fMRI segments. Panel C and D show the values of latent variables in the primary cluster and secondary cluster, respectively. The remaining 4 clusters were not shown.

## 4 Discussions

### 4.1 Innovativeness of the method

We demonstrated a new method to study the intrinsic features in resting-state fMRI using a convolutional variational autoencoder. This particular architecture has never been used to characterize rs-fMRI, although there have been a few applications in other fields that have similar convolutional VAE architectures. For example, (Kulkarni et al., 2015) used a 2-D convolutional variational autoencoder to learn intrinsic spatial patterns from images. The features of the network architecture that we developed (namely the autoencoder design, the variational approach and the 1-D convolutional layers) have many advantages for studying rs-fMRI temporal dynamics.

Firstly, the proposed method provides a parsimonious representation of brain activity by condensing it into a few highly representative components, without losing too much information. Each “brain state”, which is the collection of activity across the entire brain at any given time, can be represented as a point in a hyperplane. This brain state representation tends to be very high-dimensional. For example, the 2mm volumetric HCP data has 91×109×91=902,629 voxels, and even the greatly downsampled data examined here after parcellation with the BN atlas has 246 parcels. Extremely high-dimensional data is very sparse and hard to generalize, which is also known as the “curse of dimensionality” (Bellman, 1952). Thus, this high-dimensional definition of brain states, is overly complicated and redundant, because the many resting state networks are spatially organized, and the temporal dynamics involved may also be governed by certain rules. The “true” brain state vector may live in a much lower dimension space, which is what the VAE is designed to capture. The parsimonious representation of brain activity (using a 32-component vector to represent the brain state dynamics contained in a 246 parcels by 33 time points matrix) captures the most distinctive and prominent common trajectories that can serve as the building blocks to construct all possible spatiotemporal dynamics in resting state fMRI. This provides insight into both the spatial organization of the networks, and the characteristic dynamics for those networks. The variational approach (which includes random sampling and penalizing KL divergence) has forced the latent components to be nearly orthogonal to each other, which have helped to create a robust and unique decomposition of the brain activity and make the latent variables easier to interpret since they are disentangled.

Secondly, the use of 1-D convolutional layers has taken the structure along temporal dimension into consideration, which enables the method to simultaneously extract not only spatial patterns, but also their temporal dynamics. As discussed later, most analysis methods consider spatial information and temporal information independently. The learned latent representations were grouped into a few clusters that show similar spatial configurations that are in agreement with the anticorrelated DMN-TPN and principal gradients. Moreover, the temporal dynamics within these spatial configurations were also provided in the form of a few orthogonal components, where the one with the highest variance closely resembles the primary QPP, while the others show temporal structures that were previously extensively reported in the past.

Thirdly, as a deep learning method, this method makes minimal assumptions about rs-fMRI. The use of 1-D convolutional kernel implies that the rule governing the temporal dynamics is shift-invariant along time and is applicable to all subjects, which is a reasonable assumption to make if the goal is to find common spatiotemporal features that exist across subjects. Other than that, the neural network itself does not make any other assumptions about rs-fMRI. It is a data-driven approach that aims to extract the most common and most prominent features existing in rs-fMRI with no prior knowledge that could potentially bias the results.

### 4.2 Comparison with existing methods

While there is no existing method that strictly focuses on the same goal as the proposed method, many other methods are conceptually related, and the spatial configurations obtained by the proposed method can be compared with existing methods. In this section we compare the results from our VAE approach to other existing methods for rs-fMRI analysis, including principal component analysis (PCA), principal gradients, QPP, SWC, ICA, CAP and HMM.

#### 4.2.1 Relation to principal component analysis (PCA)

PCA and the VAE used for this study share some similarities. Both methods identify orthogonal bases for the original data and can achieve dimensionality reduction by selecting a few components that explain the most variance. In fact, a two-layer VAE with a linear activation function produces almost identical results to PCA, because both methods aim to create a linear projection of the data to an orthogonal space (Plaut, 2018). In our VAE, there are 10 layers in total, making the VAE capable of creating a much more nonlinear mapping that might capture features that would not be found in a linear mapping. On top of that, the proposed VAE has three 1-D convolutional layers to extract characteristic temporal dynamics, which are not captured by PCA. Thus the latent variables in our model captures spatiotemporal dynamics, whereas the traditional PCA often gives eigenvectors in the spatial domain, e.g. (Leonardi et al., 2013).

#### 4.2.2 Relation to diffusion embedding (principal functional connectivity gradients)

Principal functional connectivity gradients were described using a method called diffusion embedding, which nonlinearly maps brain region into an embedded hyperplane, where strongly connected points are close whereas loosely connected points are far apart (Margulies et al., 2016). The “gradients” that define the hyperplane reveal connectivity patterns over space. Like PCA, diffusion embedding provides information about connectivity geometry, but loses temporal information, whereas the proposed VAE provides a set of spatiotemporal patterns that demonstrate clusters of spatial organization while also providing information about characteristic temporal dynamics.

Because diffusion embedding and the VAE method emphasize different features of the rs-fMRI data, they are complementary to each other. The principal gradient is able to differentiate several networks along the gradient (e.g. DMN-FP-DA-VIS) whereas the variational autoencoder can only provide coarse locations (DMN and FP on one end, and DA, VA, VIS, SM on the other end). The VAE however, is also capable of showing temporal features and considers both the dynamics involved in brain activity and the interactions among brain regions, which the principle gradient lacks. Thus, they bring insights into different aspects of the same complicated brain system. The fact that the first two clusters of latent variables in VAE and the first two principal gradients show a consistent DMN versus TPN along the primary axis, and VIS versus SM, as well as DA versus VA along the secondary axis, is a reassuring indication of the consistency of the two approaches.

#### 4.2.3 Relation to QPP regression

The spatiotemporal feature that would activate latent variable 1 is very similar to the spatiotemporal patterns found in the primary QPP, specifically a sinusoidal wave-like fluctuation showing anti-correlation between the DMN and TPN, as shown in figure 4. The QPP picks up the most prominent feature in rs-fMRI because it iteratively averages the time points that have the highest correlation with the template to update the template, so it makes sense for such a feature to capture the largest portion of the variance. Aside from the primary QPP, a set of secondary QPPs have been obtained from mouse (Belloy et al., 2018) and human (Yousefi and Keilholz, 2020) resting-state fMRI data, by recursively regressing out QPP components. QPP regression is similar to the VAE method in that they both extract components that are independent to each other, and they both capture reoccurring spatiotemporal patterns. However, QPP regression was often done only for the first few components, without an exhaustive search for all possible components, perhaps because of the decreased robustness involved in the recursive convolution and regression, as well as the increased computational cost. The VAE method, however, gives an overview of all spatiotemporal patterns at the same time.

#### 4.2.4 Relation to sliding window correlation (SWC)

The proposed VAE was trained with short rs-fMRI segments of approximately 24 seconds in length. Although during training the rs-fMRI segments were shuffled, during testing (shown in figure 6) there was no shuffling, and the rs-fMRI time course was essentially transformed into latent representations using a 24-second, 50%-overlapping sliding windows, in a manner similar to the sliding window correlation method. However, for the VAE approach, the windows are used to train 32 latent variables which capture the spatiotemporal dynamics, while for sliding window correlation, dynamics are represented by the time varying correlation values. K-means clustering is applied for both approaches. For the VAE, clustering is used to group latent variables by their spatial similarity. For SWC, however, clustering the time-varying correlation is the basis for “brain states” that can be defined for each time window in the scan. Since the VAE requires components to be nearly independent of each other, the resulting clusters are more unique and clearly defined, whereas in SWC, the clusters seem to have more ambiguities because different components can mix and the long window (typically around 1~2 minutes) used for correlation can obscure short-term dynamics. For example, in (Allen et al., 2014) it was shown that the brain exhibits 7 states with connectivity patterns using a SWC method, among which states 2-7 all show notable anticorrelation between default-mode regions and sensory systems, with some variations (e.g.,. state 5 and 6 separates posterior DM nodes (precuneus and PCC) from anterior and lateral parietal regions; state 6 and 7 shows positive correlation between DM and SM area, and negative correlation between SM and VIS regions). These effects manifest as a slight deviation from the average functional connectivity, whereas in our VAE method, such separations are much more clearly defined, e.g. SM versus VIS in cluster 2, and posterior DM regions versus anterior and lateral parietal DM regions in cluster 2 and cluster 4.

#### 4.2.5 Relation to independent component analysis (ICA)

The proposed method also has some similarities to ICA, another popular method for dimensionality reduction. Though both methods try to decompose the rs-fMRI signal into independent components, the approaches they take are different. ICA can be used to discover either spatially or temporally independent components. Most rs-fMRI studies use a spatial ICA (sICA) approach to find spatial components that are maximally independent in space (Calhoun et al., 2009). It is typically applied as one step in the preprocessing to create a “functional parcellation”, which is also known as intrinsic connectivity networks (ICNs), and is often applied in conjunction with further analysis methods like SWC, e.g. (Allen et al., 2014). ICA seeks to create a matrix decomposition of the entire rs-fMRI dataset, where one matrix represents spatially independent networks and the other represent the time courses of the signals from different sources. The proposed VAE, on the other hand, is trying to find characteristic spatiotemporal patterns that are independent from each other, on a much shorter time scale. It identifies instantaneous brain trajectories within a short time window (~20 seconds) that are very characteristic, so that all the dynamics in rs-fMRI can be explained by the same set of common trajectories. The counterpart of ICA’s role of creating parcellation in this study was achieved by using the Brainnetome atlas 246-region parcellation (an anatomical parcellation), which was then organized using Yeo’s 7network7-network model.

#### 4.2.6 Relation to HMM and CAP

HMM and CAP methods are explicitly designed to characterize changes in the rs-fMRI signal over time and emphasize individual time frames in the rs-fMRI time series. The VAE, on the other hand, focuses on the dynamic patterns within rs-fMRI segments, linking spatial patterns with temporal variation. Nevertheless, the spatial patterns obtained with the three methods can be compared. For the HMM method it was reported that every fMRI frame can be classified into one of the 12 brain states, which are organized into 2 metastates (Vidaurre et al., 2017). The first metastate (state 1-4), is composed of sensory (somatic, visual, and auditory) and motor regions, and the second metastate (state 6-12) covers higher order cognitive regions that include the DMN, language, and prefrontal areas. Individual states may show specific network patterns, e.g. state 4 (visual), state 6 (DMN), state 9 (Language). In the CAP method, the few frames with posterior cingulate cortex (PCC) activation (whose correlation map resembles DMN) can be decomposed into 8 different spatial patterns (Liu and Duyn, 2013). In the first 4 components, CAP1 and CAP2 more closely resemble DMN than CAP3 and CAP4, with CAP1 extending more dorsally and CAP2 more ventrally. CAP3 highlights the middle frontal gyrus (MFG, lies in FP network in Yeo’s parcellation), whereas CAP4 highlights the superior frontal gyrus (SFG, DM network) and the parahippocampus gyrus (PHG, LIM network). There are also another 4 CAPs with less resemblance to DMN that have lower within-group similarity. These variations of spatial patterns observed in the individual time frames, could emerge from the superposition of the orthogonal components found in the proposed VAE model. For example, positive components in latent cluster 1 (DMN activation) superimposed with positive components in latent cluster 2 (DM and LIM activation) could give rise to a spatial pattern similar to CAP4, whereas positive components in latent cluster 1 superimposed with negative components in latent cluster 2 (FP activation) could result in something more similar to CAP3.

### 4.3 New findings from the VAE

Although fundamentally different from existing methods, the trained VAE returns results in line with many previous studies. In particular, the DMN-TPN contrast seen here was also reported in DMN-TPN anticorrelation, QPPs, metastates (Vidaurre et al., 2017) and principal gradients. In addition to recapitulating previous findings, the VAE method also reveals some spatiotemporal trajectories that were not previously discovered or extensively discussed. For example, there are spatiotemporal trajectories that generally follow the DMN-TPN spatial configuration, but have much faster frequencies when compared to the QPP (e.g. latent variable 2, 4, 5 in the primary latent variable cluster). A second example is given by the spatiotemporal trajectories in the secondary cluster, which show temporal dynamics along a spatial distribution similar to principal gradient 2 (VIS, DA on one side and SM, VA on the other). These additional spatiotemporal dynamics are worth investigating in the future, including but not limited to their reproducibility and whether they would change under different cognitive states.

Another interesting feature to notice is that there seem to be two modes of activity revealed by the spatiotemporal trajectories. One mode has distinct on and off blocks showing two networks having exactly opposite phase (e.g. latent variable 1). Another mode shows a gradual change of phase/peak time along the spatial dimension, which behaves more like a wave propagating through different networks, which could also be related the findings in (Gu et al., 2020) and (Yousefi and Keilholz, 2020). This propagation/time lag, and how it interacts with the first mode (the well-known DMN-TPN anticorrelation) is worth investigating in the future.

### 4.4 Potential applications

The proposed VAE has found a set of characteristic spatiotemporal brain trajectories that can explain most of the dynamics involved in rs-fMRI. This new perspective provides insights into the brain’s spatiotemporal dynamics that cannot be accessed from traditional methods such as functional connectivity. Future work should explore how these characteristic trajectories change when the cognitive state is changed (e.g. task performance, sleeping vs resting state) or with the presence of a neurological or psychiatric disorder (e.g. Alzheimer’s disease, major depression disorder or ADHD) as these alterations may change the fMRI characteristics and instantaneous dynamics. For example, in (Jones et al., 2012b) it was reported that the differences in static connectivity observed in Alzheimer’s disease can be explained by differences in dwell time in DMN sub-network configurations, which suggests the dynamics of brain activity, and presumably the characteristic spatiotemporal brain trajectories are also altered by Alzheimer’s disease. Potential additional approaches include training multiple VAEs on patient and control groups to see if the characteristic trajectories identified are different, using the characteristic trajectories from healthy resting-state data as a benchmark to identify statistical differences among groups, or training a classifier to utilize trajectories to identify the cognitive state or neurological disorder.

### 4.5 Technical limitations

Although the proposed method has opened up a new perspective for viewing rs-fMRI dynamics, it does have some technical limitations. First many of the hyperparameters are empirically chosen, which is almost always not the most “optimal” solution of the problem. While it is possible to perform an exhaustive grid search for optimal parameters in some circumstances, the computational cost quickly become infeasible when the number and the range of parameters being tuned is large (Wu et al., 2019). That said, we did consider many factors when designing the neural network so that the parameters involved are within a reasonable range. For example, the number of layers cannot be too small, or the model will lack expressive power and cannot capture complicated features; on the other hand, the number of layers cannot be too large or the gradient will not backpropagate easily, resulting in difficulties in training. We also performed a holdout validation to examine the effect of the hyperparameters like the number of latent variables and number of layers (the results were shown in supplementary materials section S.1). While the choices we made are not necessarily the best, they certainly are not the worst.

Secondly the network was trained with a built-in “sliding window”. We chose to divide the dataset into 50% overlapping, 33-TR (24 sec) long time window. This is likely to limit the lowest frequency component the model can identify, which is around 1/24 = 0.042Hz. Fluctuations that occur at lower frequencies are likely to be ignored by the model. Using a longer window may help capture components with lower frequencies, but doing such also requires an increase number of latent variables to encode the additional information in the elongated window, thus making the model more complex and harder to train. Eventually there will be a soft limit of how long the window can be feasibly implemented, which puts a lower bound to the frequencies that can be properly identified.

Thirdly the proposed method is tailored for parcellated rs-fMRI data. For nonparcellated rs-fMRI data, it would make more sense to use multidimensional convolutional layers instead of the 1-D convolutional layers we used in our work, since volumetric rs-fMRI data may preserve the property of shift-invariance not only in the time domain, but also in the spatial domain as well. However, volumetric rs-fMRI data are orders of magnitude larger than parcellated rs-fMRI data in size, whose modeling demands a neural network with more complex structure and greater expressive power. This increased model size makes the network harder to train. Whether it is possible to train such a model for nonparcellated rs-fMRI data, and if so how the latent variable would differ from those obtained from a model trained with parcellated rs-fMRI data, still remains unknown at the moment.

## 5 Conclusion

In this article we proposed a novel convolutional variational autoencoder to extract intrinsic spatiotemporal patterns from short segments of resting-state fMRI data. The extracted latent dimensions show clear clusters in the spatial domain that are in agreement with previous findings, but also provide temporal information about the evolution of brain activity as well. Some spatiotemporal features were similar to previously-described QPPs, but there are others with smaller variances that were not previously discovered, which is worth investigating in the future.

## Acknowledgements

Funding sources: NIH 1 R01NS078095-01, BRAIN initiative R01 MH 111416 and NSF INSPIRE. The authors would like to thank the Washington University–University of Minnesota Consortium of the Human Connectome Project (WU-Minn HCP) for generating and making publicly available the HCP data. The authors would like to thank Chinese Scholarship Council (CSC) for financial support.

