## supplemental materials for "Spatiotemporal Trajectories in Resting-state FMRI Revealed by Convolutional Variational Autoencoder"

#### S1. Evaluation of different network architectures.

To evaluate the performance of different network architectures, we trained 5 different VAEs using the same procedure. The networks differ either in width (number of hidden units in the final encoder layer, or the number of latent variables), or depth (total number of layers). The validation loss of each VAE is shown in Figure S1. It can be seen that model 1 (narrow) and model 5 (deep) has significantly higher validation loss.

Model 1 has higher validation loss because it is too narrow in the latent layer, thus the bottleneck effect is too strong so that the model lacks expressive power to capture the details of the signal and the reconstruction becomes suboptimal. Model 5 has higher validation loss because it is too deep, thus the gradient cannot backpropagate easily. Training such deeper networks may require residual connections, which we did not explore in this paper.

The rest 3 models have comparable performance in terms of validation loss, however the spatiotemporal trajectories exhibited by the latent variables are different, which is how we determined the best model for our study. Model 3 has 64 latent variables, but only 34 of them exhibit meaningful patterns, whereas the remaining 30 were nulled during the training during the training, suggesting that they are redundant. The nulling effect can be seen in figure S3, where the latent variables exhibit a constant spatial pattern regardless of the change in the latent variables, and have zero variance explained. This is caused by the regularizing effect of the K-L divergence, which encourages sparsity in the latent space. Thus the model is unnecessarily wide and we chose our model to have 32 latent variables in the end.

**Table S1. Network architectures compared in the evaluation.** In this table, the output size for each layer were shown next to the name of the corresponding layer (for CONV layers the size is expressed as number of channels x number of time points). The number of hidden units (latent variables) is equal to the output size of the final FC layer. The number of layer for each model is twice the size of the number of layers in the encoder (since the decoder is symmetric to the encoder). Compared to the final selected model (model 2), the rest of the models either differ in width (number of hidden units in the final encoder layer) or in depth (number of layers).

| Model | Model 1 (narrow)<br>16-unit, 10-layer | **Model 2 (selected)<br>32-unit, 10-layer** | Model 3 (wide)<br>64-unit, 10-layer | Model 4 (shallow)<br>32-unit, 4-layer | Model 5 (deep)<br>32-unit, 16-layer |
|---|---|---|---|---|---|
| Encoder Structure | CONV1 128X17<br>CONV2 64X9<br>CONV3 64X6<br>FC1 256<br>FC2 16 | CONV1 128X17<br>CONV2 64X9<br>CONV3 64X6<br>FC1 256<br>FC2 32 | CONV1 128X17<br>CONV2 64X9<br>CONV3 64X6<br>FC1 256<br>FC2 64 | FC1 256<br>FC2 32 | CONV1 256X17<br>CONV2 128X17<br>CONV3 128X9<br>CONV3 64X9<br>CONV3 64X6<br>FC1 256<br>FC2 128<br>FC3 32 |
| Decoder Structure | Symmetric to Encoder | | | | |

CONV, convolutional layer; FC, fully-connected layer

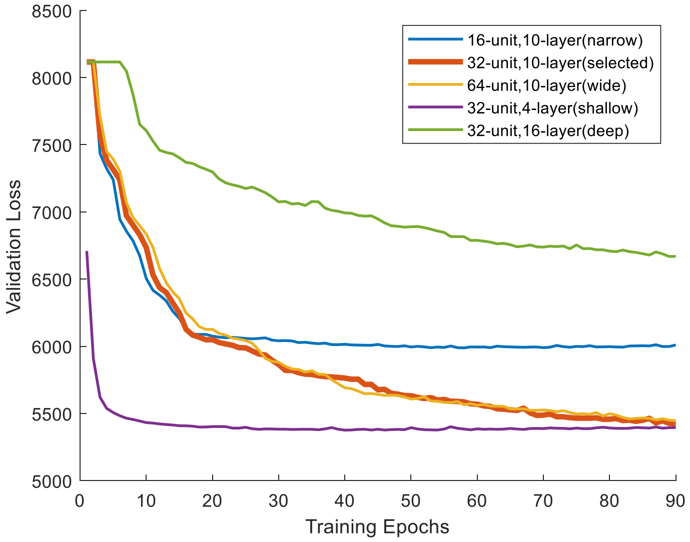


**Figure S2. Validation loss of the 5 models during training.**

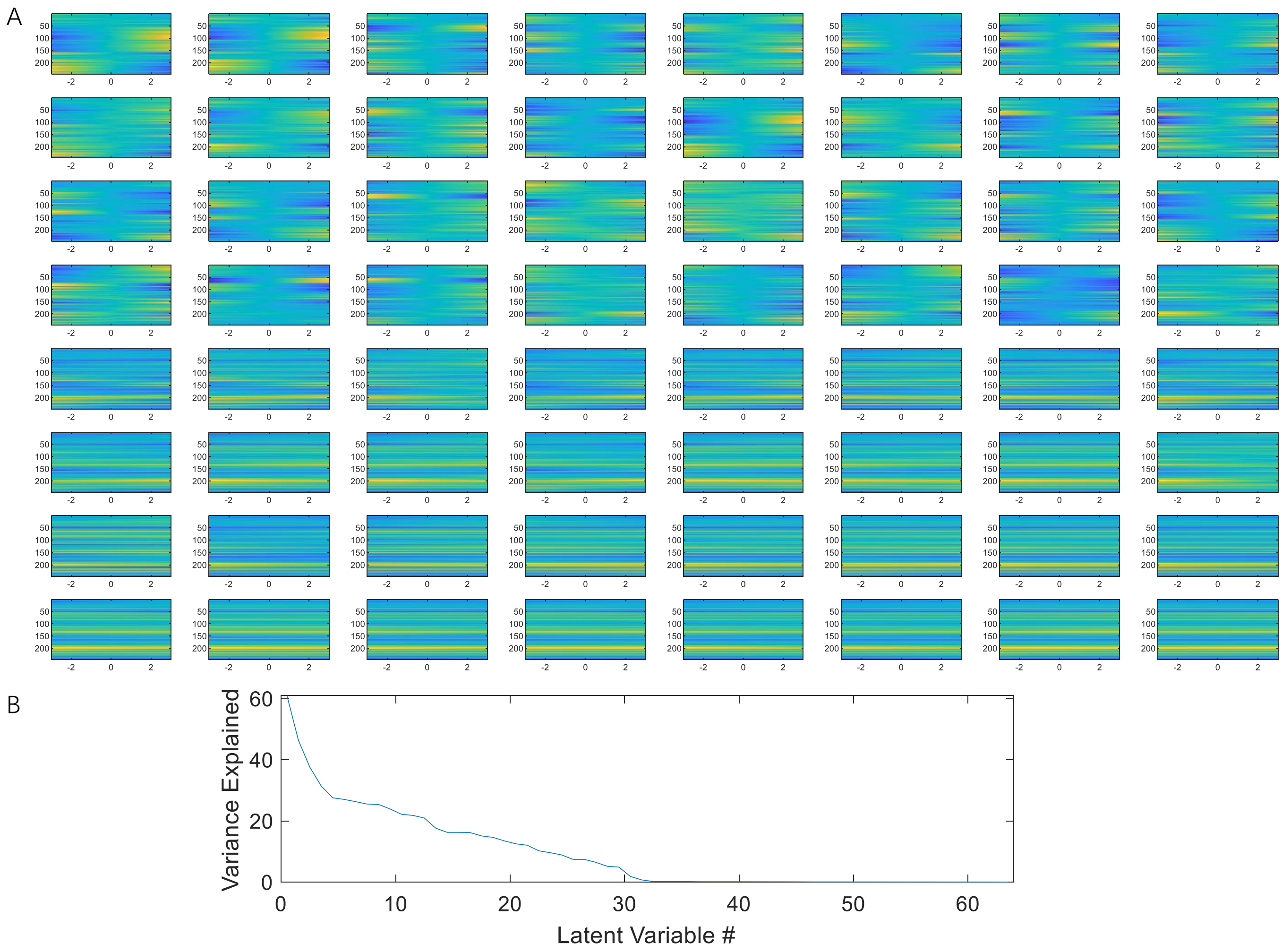


**Figure S3. The spatial profile (A) and the variance explained (B) of the 64 latent variables obtained by model 3 (wide model).** The spatial profile at the max-variance time was shown as a function of the corresponding latent variable sliding from -3 to +3, in a manner similar to figure 2 panel A. It can be seen that latent variable 35-64 (in total 30 latent variables) all exhibit a constant spatial profile that is not changing along the latent variables, suggests they don't contribute to the encoding and decoding. Their variances explained is thus also zero.

Model 4 has only 2 layers in either encoder and decoder, thus has the simplest form of nonlinearity. Despite it shows slightly lower validation loss (meaning more accurate reconstruction), its latent features are much less organized, potentially due to the lack of expressive power required for extracting features with complicated nonlinearity. It can be seen in figure S4 that the clustering of latent variables based on spatial profiles (in a manner similar to section 3.2, figure 3) results in much blurred boundaries among latent variable clusters compared to the selected 10-layer model presented in the paper. The variance explained is also more evenly spread out across all 32 latent variables, whereas in the 10-layer model, the first few clusters clearly show much higher variance explained than the rest. This suggests that the 10-layer model we used in this study has much better expressive power than the shallower counterpart, as a result the latent variables it found were much more representative and characteristic.

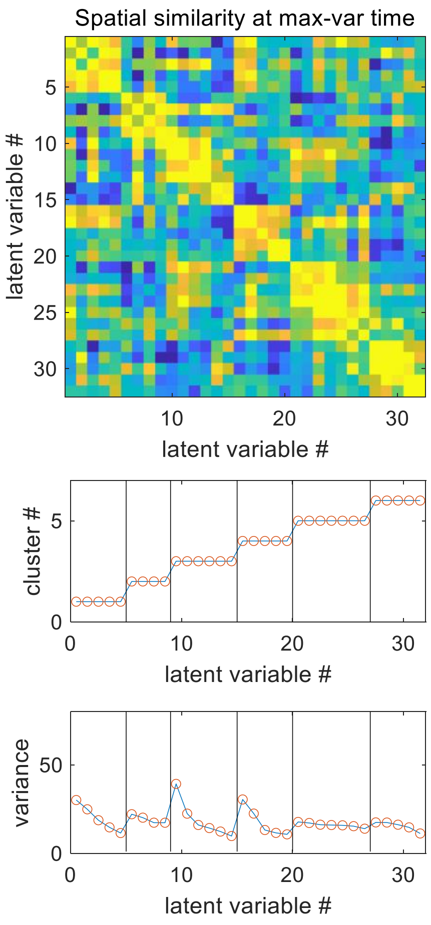


**Figure S4. The latent variables obtained by model 4 (shallow) can also be clustered using K means clustering (K = 6) based on spatial similarity, but they exhibit much more blurred boundaries among clusters compared to model 2 (selected) shown in figure 3.** The spatial similarities among latent variables at the max-variance time, measured by Pearson correlation between the spatial profiles. The cluster label index and the variance explained are also shown.

### S2. Compare the orthogonality of latent variables for networks trained with beta =1 and beta=4

The orthogonality of latent variables is an important feature because it guarantees a unique solution and thus improves the model's robustness and reproducibility. It also makes different latent variables disentangled and thus help with interpreting the results. Despite theoretically penalizing K-L divergence will give latent variables nearly orthogonal to each other, it is worthwhile to validate the orthogonality of the beta-VAE model, and compare with the plain VAE model.

To do that, we calculated the Pearson correlation among the spatiotemporal patterns of the latent variables, as well as the Pearson correlation among the latent variable time courses. The spatiotemporal patterns of the latent variables are like the bases of the hyperplane where brain state exists, whereas the latent variable time courses show the projection of the brain state on these bases at any given time. The spatiotemporal patterns are 2D matrices, so in order to calculate the correlation among them, they were flattened. Ideally if the latent variables are truly orthogonal to each other, both correlation matrices will be an identity matrix. In Figure S5 it can be seen that both correlation matrices of the beta-VAE model are closer to an identity matrix than the plain-VAE model, suggesting that it is more robust and regularized as intended.

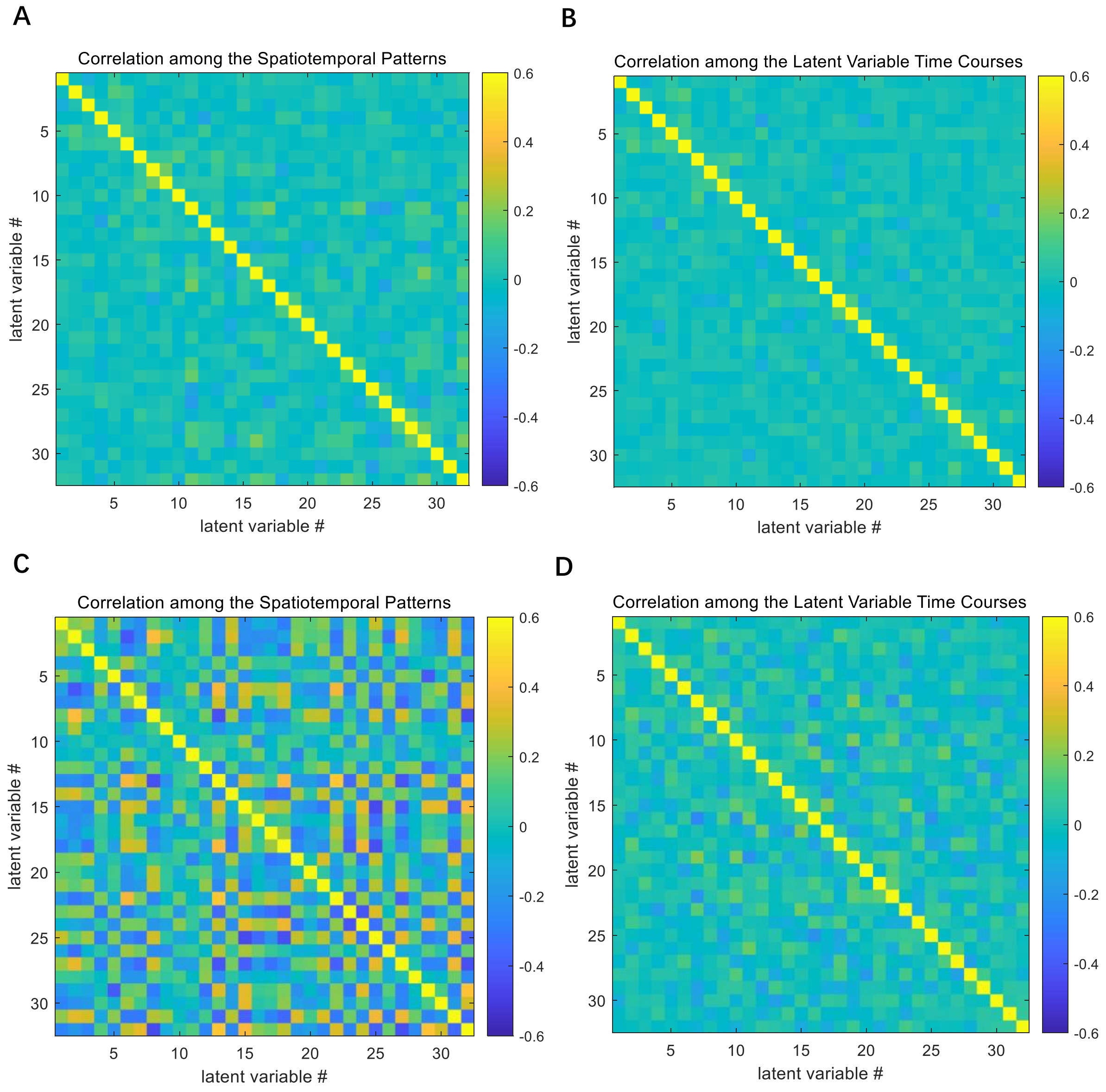


**Figure S5. The latent variables in beta-VAE (beta = 4) are more orthogonal than plain-VAE (beta = 1).** Panel A, correlation among the spatiotemporal patterns obtained by beta-VAE model. Panel B, correlation among the latent variable time courses obtained by beta-VAE model. Panel C, correlation among the spatiotemporal patterns obtained by plain-VAE model. Panel D, correlation among the latent variable time courses obtained by plain-VAE model.

### S3. Location of the 246 parcels

| | Parcel ID | BN parcel ID | Lobe | Gyrus | Region | Subregion Name |
|---|---|---|---|---|---|---|
| Subcortical Regions (SC) | 1 | 165 | Insular Lobe | Insular Gyrus | INS_L_6_2 | vIa |
| | 2 | 177 | Limbic Lobe | Cingulate Gyrus | CG_L_7_2 | A24rv |
| | 3 | 178 | Limbic Lobe | Cingulate Gyrus | CG_R_7_2 | A24rv |
| | 4 | 211 | Subcortical Nuclei | Amygdala | Amyg_L_2_1 | mAmyg |
| | 5 | 212 | Subcortical Nuclei | Amygdala | Amyg_R_2_1 | mAmyg |
| | 6 | 213 | Subcortical Nuclei | Amygdala | Amyg_L_2_2 | lAmyg |
| | 7 | 214 | Subcortical Nuclei | Amygdala | Amyg_R_2_2 | lAmyg |
| | 8 | 215 | Subcortical Nuclei | Hippocampus | Hipp_L_2_1 | rHipp |
| | 9 | 216 | Subcortical Nuclei | Hippocampus | Hipp_R_2_1 | rHipp |
| | 10 | 217 | Subcortical Nuclei | Hippocampus | Hipp_L_2_2 | cHipp |
| | 11 | 218 | Subcortical Nuclei | Hippocampus | Hipp_R_2_2 | cHipp |
| | 12 | 219 | Subcortical Nuclei | Basal Ganglia | BG_L_6_1 | vCa |
| | 13 | 220 | Subcortical Nuclei | Basal Ganglia | BG_R_6_1 | vCa |
| | 14 | 221 | Subcortical Nuclei | Basal Ganglia | BG_L_6_2 | GP |
| | 15 | 222 | Subcortical Nuclei | Basal Ganglia | BG_R_6_2 | GP |
| | 16 | 223 | Subcortical Nuclei | Basal Ganglia | BG_L_6_3 | NAC |
| | 17 | 224 | Subcortical Nuclei | Basal Ganglia | BG_R_6_3 | NAC |
| | 18 | 225 | Subcortical Nuclei | Basal Ganglia | BG_L_6_4 | vmPu |
| | 19 | 226 | Subcortical Nuclei | Basal Ganglia | BG_R_6_4 | vmPu |
| | 20 | 227 | Subcortical Nuclei | Basal Ganglia | BG_L_6_5 | dCa |
| | 21 | 228 | Subcortical Nuclei | Basal Ganglia | BG_R_6_5 | dCa |
| | 22 | 229 | Subcortical Nuclei | Basal Ganglia | BG_L_6_6 | dlPu |
| | 23 | 230 | Subcortical Nuclei | Basal Ganglia | BG_R_6_6 | dlPu |
| | 24 | 231 | Subcortical Nuclei | Thalamus | Tha_L_8_1 | mPFtha |
| | 25 | 232 | Subcortical Nuclei | Thalamus | Tha_R_8_1 | mPFtha |
| | 26 | 233 | Subcortical Nuclei | Thalamus | Tha_L_8_2 | mPMtha |
| | 27 | 234 | Subcortical Nuclei | Thalamus | Tha_R_8_2 | mPMtha |
| | 28 | 235 | Subcortical Nuclei | Thalamus | Tha_L_8_3 | Stha |
| | 29 | 236 | Subcortical Nuclei | Thalamus | Tha_R_8_3 | Stha |
| | 30 | 237 | Subcortical Nuclei | Thalamus | Tha_L_8_4 | rTtha |
| | 31 | 238 | Subcortical Nuclei | Thalamus | Tha_R_8_4 | rTtha |
| | 32 | 239 | Subcortical Nuclei | Thalamus | Tha_L_8_5 | PPtha |
| | 33 | 240 | Subcortical Nuclei | Thalamus | Tha_R_8_5 | PPtha |
| | 34 | 241 | Subcortical Nuclei | Thalamus | Tha_L_8_6 | Otha |
| | 35 | 242 | Subcortical Nuclei | Thalamus | Tha_R_8_6 | Otha |
| | 36 | 243 | Subcortical Nuclei | Thalamus | Tha_L_8_7 | cTtha |
| | 37 | 244 | Subcortical Nuclei | Thalamus | Tha_R_8_7 | cTtha |
| | 38 | 245 | Subcortical Nuclei | Thalamus | Tha_L_8_8 | lPFtha |
| | 39 | 246 | Subcortical Nuclei | Thalamus | Tha_R_8_8 | lPFtha |
| Visual (VIS) | 40 | 105 | Temporal Lobe | Fusiform Gyrus | FuG_L_3_2 | A37mv |
| | 41 | 106 | Temporal Lobe | Fusiform Gyrus | FuG_R_3_2 | A37mv |
| | 42 | 108 | Temporal Lobe | Fusiform Gyrus | FuG_R_3_3 | A37lv |
| | 43 | 112 | Temporal Lobe | Parahippocampal Gyrus | PhG_R_6_2 | A35/36c |
| | 44 | 113 | Temporal Lobe | Parahippocampal Gyrus | PhG_L_6_3 | TL |
| | 45 | 114 | Temporal Lobe | Parahippocampal Gyrus | PhG_R_6_3 | TL |
| | 46 | 119 | Temporal Lobe | Parahippocampal Gyrus | PhG_L_6_6 | TH |
| | 47 | 120 | Temporal Lobe | Parahippocampal Gyrus | PhG_R_6_6 | TH |
| | 48 | 135 | Parietal Lobe | Inferior Parietal Lobule | IPL_L_6_1 | A39c |
| | 49 | 136 | Parietal Lobe | Inferior Parietal Lobule | IPL_R_6_1 | A39c |
| | 50 | 151 | Parietal Lobe | Precuneus | PCun_L_4_3 | dmPOS |
| | 51 | 152 | Parietal Lobe | Precuneus | PCun_R_4_3 | dmPOS |
| | 52 | 182 | Limbic Lobe | Cingulate Gyrus | CG_R_7_4 | A23v |
| | 53 | 189 | Occipital Lobe | MedioVentral Occipital Cortex | MVOcC_L_5_1 | cLinG |
| | 54 | 190 | Occipital Lobe | MedioVentral Occipital Cortex | MVOcC_R_5_1 | cLinG |
| | 55 | 191 | Occipital Lobe | MedioVentral Occipital Cortex | MVOcC_L_5_2 | rCunG |
| | 56 | 192 | Occipital Lobe | MedioVentral Occipital Cortex | MVOcC_R_5_2 | rCunG |

| | | | | | | |
|---|---|---|---|---|---|---|
| | 57 | 193 | Occipital Lobe | MedioVentral Occipital Cortex | MVOcC_L_5_3 | cCunG |
| | 58 | 194 | Occipital Lobe | MedioVentral Occipital Cortex | MVOcC_R_5_3 | cCunG |
| | 59 | 195 | Occipital Lobe | MedioVentral Occipital Cortex | MVOcC_L_5_4 | rLinG |
| | 60 | 196 | Occipital Lobe | MedioVentral Occipital Cortex | MVOcC_R_5_4 | rLinG |
| | 61 | 197 | Occipital Lobe | MedioVentral Occipital Cortex | MVOcC_L_5_5 | vmPOS |
| | 62 | 198 | Occipital Lobe | MedioVentral Occipital Cortex | MVOcC_R_5_5 | vmPOS |
| | 63 | 199 | Occipital Lobe | Lateral Occipital Cortex | LOcC_L_4_1 | mOccG |
| | 64 | 200 | Occipital Lobe | Lateral Occipital Cortex | LOcC_R_4_1 | mOccG |
| | 65 | 202 | Occipital Lobe | Lateral Occipital Cortex | LOcC_R_4_2 | V5/MT+ |
| | 66 | 203 | Occipital Lobe | Lateral Occipital Cortex | LOcC_L_4_3 | OPC |
| | 67 | 204 | Occipital Lobe | Lateral Occipital Cortex | LOcC_R_4_3 | OPC |
| | 68 | 205 | Occipital Lobe | Lateral Occipital Cortex | LOcC_L_4_4 | iOccG |
| | 69 | 206 | Occipital Lobe | Lateral Occipital Cortex | LOcC_R_4_4 | iOccG |
| | 70 | 207 | Occipital Lobe | Lateral Occipital Cortex | LOcC_L_2_1 | msOccG |
| | 71 | 208 | Occipital Lobe | Lateral Occipital Cortex | LOcC_R_2_1 | msOccG |
| | 72 | 209 | Occipital Lobe | Lateral Occipital Cortex | LOcC_L_2_2 | lsOccG |
| | 73 | 210 | Occipital Lobe | Lateral Occipital Cortex | LOcC_R_2_2 | lsOccG |
| Somatomotor (SM) | 74 | 9 | Frontal Lobe | Superior Frontal Gyrus | SFG_L_7_5 | A6m |
| | 75 | 10 | Frontal Lobe | Superior Frontal Gyrus | SFG_R_7_5 | A6m |
| | 76 | 53 | Frontal Lobe | Precentral Gyrus | PrG_L_6_1 | A4hf |
| | 77 | 54 | Frontal Lobe | Precentral Gyrus | PrG_R_6_1 | A4hf |
| | 78 | 57 | Frontal Lobe | Precentral Gyrus | PrG_L_6_3 | A4ul |
| | 79 | 58 | Frontal Lobe | Precentral Gyrus | PrG_R_6_3 | A4ul |
| | 80 | 59 | Frontal Lobe | Precentral Gyrus | PrG_L_6_4 | A4t |
| | 81 | 60 | Frontal Lobe | Precentral Gyrus | PrG_R_6_4 | A4t |
| | 82 | 66 | Frontal Lobe | Paracentral Lobule | PCL_R_2_1 | A1/2/3ll |
| | 83 | 67 | Frontal Lobe | Paracentral Lobule | PCL_L_2_2 | A4ll |
| | 84 | 68 | Frontal Lobe | Paracentral Lobule | PCL_R_2_2 | A4ll |
| | 85 | 71 | Temporal Lobe | Superior Temporal Gyrus | STG_L_6_2 | A41/42 |
| | 86 | 72 | Temporal Lobe | Superior Temporal Gyrus | STG_R_6_2 | A41/42 |
| | 87 | 73 | Temporal Lobe | Superior Temporal Gyrus | STG_L_6_3 | TE1.0/TE1.2 |
| | 88 | 74 | Temporal Lobe | Superior Temporal Gyrus | STG_R_6_3 | TE1.0/TE1.2 |
| | 89 | 75 | Temporal Lobe | Superior Temporal Gyrus | STG_L_6_4 | A22c |
| | 90 | 76 | Temporal Lobe | Superior Temporal Gyrus | STG_R_6_4 | A22c |
| | 91 | 131 | Parietal Lobe | Superior Parietal Lobule | SPL_L_5_4 | A7pc |
| | 92 | 132 | Parietal Lobe | Superior Parietal Lobule | SPL_R_5_4 | A7pc |
| | 93 | 145 | Parietal Lobe | Inferior Parietal Lobule | IPL_L_6_6 | A40rv |
| | 94 | 146 | Parietal Lobe | Inferior Parietal Lobule | IPL_R_6_6 | A40rv |
| | 95 | 149 | Parietal Lobe | Precuneus | PCun_L_4_2 | A5m |
| | 96 | 155 | Parietal Lobe | Postcentral Gyrus | PoG_L_4_1 | A1/2/3ulhf |
| | 97 | 156 | Parietal Lobe | Postcentral Gyrus | PoG_R_4_1 | A1/2/3ulhf |
| | 98 | 157 | Parietal Lobe | Postcentral Gyrus | PoG_L_4_2 | A1/2/3tonIa |
| | 99 | 158 | Parietal Lobe | Postcentral Gyrus | PoG_R_4_2 | A1/2/3tonIa |
| | 100 | 160 | Parietal Lobe | Postcentral Gyrus | PoG_R_4_3 | A2 |
| | 101 | 161 | Parietal Lobe | Postcentral Gyrus | PoG_L_4_4 | A1/2/3tru |
| | 102 | 162 | Parietal Lobe | Postcentral Gyrus | PoG_R_4_4 | A1/2/3tru |
| | 103 | 163 | Insular Lobe | Insular Gyrus | INS_L_6_1 | G |
| | 104 | 164 | Insular Lobe | Insular Gyrus | INS_R_6_1 | G |
| | 105 | 171 | Insular Lobe | Insular Gyrus | INS_L_6_5 | dIg |
| | 106 | 172 | Insular Lobe | Insular Gyrus | INS_R_6_5 | dIg |
| Dorsal Attention (DA) | 107 | 7 | Frontal Lobe | Superior Frontal Gyrus | SFG_L_7_4 | A6dl |
| | 108 | 8 | Frontal Lobe | Superior Frontal Gyrus | SFG_R_7_4 | A6dl |
| | 109 | 25 | Frontal Lobe | Middle Frontal Gyrus | MFG_L_7_6 | A6vl |
| | 110 | 26 | Frontal Lobe | Middle Frontal Gyrus | MFG_R_7_6 | A6vl |
| | 111 | 30 | Frontal Lobe | Inferior Frontal Gyrus | IFG_R_6_1 | A44d |
| | 112 | 55 | Frontal Lobe | Precentral Gyrus | PrG_L_6_2 | A6cdl |
| | 113 | 56 | Frontal Lobe | Precentral Gyrus | PrG_R_6_2 | A6cdl |
| | 114 | 63 | Frontal Lobe | Precentral Gyrus | PrG_L_6_6 | A6cvl |
| | 115 | 64 | Frontal Lobe | Precentral Gyrus | PrG_R_6_6 | A6cvl |
| | 116 | 85 | Temporal Lobe | Middle Temporal Gyrus | MTG_L_4_3 | A37dl |

| | | | | | | |
|---|---|---|---|---|---|---|
| | 117 | 86 | Temporal Lobe | Middle Temporal Gyrus | MTG_R_4_3 | A37dl |
| | 118 | 91 | Temporal Lobe | Inferior Temporal Gyrus | ITG_L_7_2 | A37elv |
| | 119 | 92 | Temporal Lobe | Inferior Temporal Gyrus | ITG_R_7_2 | A37elv |
| | 120 | 97 | Temporal Lobe | Inferior Temporal Gyrus | ITG_L_7_5 | A37vl |
| | 121 | 98 | Temporal Lobe | Inferior Temporal Gyrus | ITG_R_7_5 | A37vl |
| | 122 | 107 | Temporal Lobe | Fusiform Gyrus | FuG_L_3_3 | A37lv |
| | 123 | 125 | Parietal Lobe | Superior Parietal Lobule | SPL_L_5_1 | A7r |
| | 124 | 126 | Parietal Lobe | Superior Parietal Lobule | SPL_R_5_1 | A7r |
| | 125 | 127 | Parietal Lobe | Superior Parietal Lobule | SPL_L_5_2 | A7c |
| | 126 | 128 | Parietal Lobe | Superior Parietal Lobule | SPL_R_5_2 | A7c |
| | 127 | 129 | Parietal Lobe | Superior Parietal Lobule | SPL_L_5_3 | A5l |
| | 128 | 130 | Parietal Lobe | Superior Parietal Lobule | SPL_R_5_3 | A5l |
| | 129 | 133 | Parietal Lobe | Superior Parietal Lobule | SPL_L_5_5 | A7ip |
| | 130 | 134 | Parietal Lobe | Superior Parietal Lobule | SPL_R_5_5 | A7ip |
| | 131 | 139 | Parietal Lobe | Inferior Parietal Lobule | IPL_L_6_3 | A40rd |
| | 132 | 140 | Parietal Lobe | Inferior Parietal Lobule | IPL_R_6_3 | A40rd |
| | 133 | 143 | Parietal Lobe | Inferior Parietal Lobule | IPL_L_6_5 | A39rv |
| | 134 | 150 | Parietal Lobe | Precuneus | PCun_R_4_2 | A5m |
| | 135 | 159 | Parietal Lobe | Postcentral Gyrus | PoG_L_4_3 | A2 |
| | 136 | 201 | Occipital Lobe | Lateral Occipital Cortex | LOcC_L_4_2 | V5/MT+ |
| Ventral Attention (VA) | 137 | 2 | Frontal Lobe | Superior Frontal Gyrus | SFG_R_7_1 | A8m |
| | 138 | 15 | Frontal Lobe | Middle Frontal Gyrus | MFG_L_7_1 | A9/46d |
| | 139 | 37 | Frontal Lobe | Inferior Frontal Gyrus | IFG_L_6_5 | A44op |
| | 140 | 38 | Frontal Lobe | Inferior Frontal Gyrus | IFG_R_6_5 | A44op |
| | 141 | 39 | Frontal Lobe | Inferior Frontal Gyrus | IFG_L_6_6 | A44v |
| | 142 | 40 | Frontal Lobe | Inferior Frontal Gyrus | IFG_R_6_6 | A44v |
| | 143 | 61 | Frontal Lobe | Precentral Gyrus | PrG_L_6_5 | A4tl |
| | 144 | 62 | Frontal Lobe | Precentral Gyrus | PrG_R_6_5 | A4tl |
| | 145 | 65 | Frontal Lobe | Paracentral Lobule | PCL_L_2_1 | A1/2/3ll |
| | 146 | 123 | Temporal Lobe | Posterior Superior Temporal Sulcus | pSTS_L_2_2 | cpSTS |
| | 147 | 124 | Temporal Lobe | Posterior Superior Temporal Sulcus | pSTS_R_2_2 | cpSTS |
| | 148 | 167 | Insular Lobe | Insular Gyrus | INS_L_6_3 | dIa |
| | 149 | 168 | Insular Lobe | Insular Gyrus | INS_R_6_3 | dIa |
| | 150 | 169 | Insular Lobe | Insular Gyrus | INS_L_6_4 | vId/vIg |
| | 151 | 170 | Insular Lobe | Insular Gyrus | INS_R_6_4 | vId/vIg |
| | 152 | 173 | Insular Lobe | Insular Gyrus | INS_L_6_6 | dId |
| | 153 | 174 | Insular Lobe | Insular Gyrus | INS_R_6_6 | dId |
| | 154 | 180 | Limbic Lobe | Cingulate Gyrus | CG_R_7_3 | A32p |
| | 155 | 183 | Limbic Lobe | Cingulate Gyrus | CG_L_7_5 | A24cd |
| | 156 | 184 | Limbic Lobe | Cingulate Gyrus | CG_R_7_5 | A24cd |
| | 157 | 185 | Limbic Lobe | Cingulate Gyrus | CG_L_7_6 | A23c |
| | 158 | 186 | Limbic Lobe | Cingulate Gyrus | CG_R_7_6 | A23c |
| Limbic (LIM) | 159 | 27 | Frontal Lobe | Middle Frontal Gyrus | MFG_L_7_7 | A10l |
| | 160 | 45 | Frontal Lobe | Orbital Gyrus | OrG_L_6_3 | A11l |
| | 161 | 47 | Frontal Lobe | Orbital Gyrus | OrG_L_6_4 | A11m |
| | 162 | 48 | Frontal Lobe | Orbital Gyrus | OrG_R_6_4 | A11m |
| | 163 | 49 | Frontal Lobe | Orbital Gyrus | OrG_L_6_5 | A13 |
| | 164 | 50 | Frontal Lobe | Orbital Gyrus | OrG_R_6_5 | A13 |
| | 165 | 69 | Temporal Lobe | Superior Temporal Gyrus | STG_L_6_1 | A38m |
| | 166 | 70 | Temporal Lobe | Superior Temporal Gyrus | STG_R_6_1 | A38m |
| | 167 | 77 | Temporal Lobe | Superior Temporal Gyrus | STG_L_6_5 | A38l |
| | 168 | 78 | Temporal Lobe | Superior Temporal Gyrus | STG_R_6_5 | A38l |
| | 169 | 89 | Temporal Lobe | Inferior Temporal Gyrus | ITG_L_7_1 | A20iv |
| | 170 | 90 | Temporal Lobe | Inferior Temporal Gyrus | ITG_R_7_1 | A20iv |
| | 171 | 93 | Temporal Lobe | Inferior Temporal Gyrus | ITG_L_7_3 | A20r |
| | 172 | 94 | Temporal Lobe | Inferior Temporal Gyrus | ITG_R_7_3 | A20r |
| | 173 | 96 | Temporal Lobe | Inferior Temporal Gyrus | ITG_R_7_4 | A20il |
| | 174 | 101 | Temporal Lobe | Inferior Temporal Gyrus | ITG_L_7_7 | A20cv |
| | 175 | 102 | Temporal Lobe | Inferior Temporal Gyrus | ITG_R_7_7 | A20cv |
| | 176 | 103 | Temporal Lobe | Fusiform Gyrus | FuG_L_3_1 | A20rv |

| | | | | | | |
|---|---|---|---|---|---|---|
| | 177 | 104 | Temporal Lobe | Fusiform Gyrus | FuG_R_3_1 | A20rv |
| | 178 | 109 | Temporal Lobe | Parahippocampal Gyrus | PhG_L_6_1 | A35/36r |
| | 179 | 110 | Temporal Lobe | Parahippocampal Gyrus | PhG_R_6_1 | A35/36r |
| | 180 | 111 | Temporal Lobe | Parahippocampal Gyrus | PhG_L_6_2 | A35/36c |
| | 181 | 115 | Temporal Lobe | Parahippocampal Gyrus | PhG_L_6_4 | A28/34 |
| | 182 | 116 | Temporal Lobe | Parahippocampal Gyrus | PhG_R_6_4 | A28/34 |
| | 183 | 117 | Temporal Lobe | Parahippocampal Gyrus | PhG_L_6_5 | TI |
| | 184 | 118 | Temporal Lobe | Parahippocampal Gyrus | PhG_R_6_5 | TI |
| Frontalparietal (FP) | 185 | 1 | Frontal Lobe | Superior Frontal Gyrus | SFG_L_7_1 | A8m |
| | 186 | 4 | Frontal Lobe | Superior Frontal Gyrus | SFG_R_7_2 | A8dl |
| | 187 | 12 | Frontal Lobe | Superior Frontal Gyrus | SFG_R_7_6 | A9m |
| | 188 | 16 | Frontal Lobe | Middle Frontal Gyrus | MFG_R_7_1 | A9/46d |
| | 189 | 17 | Frontal Lobe | Middle Frontal Gyrus | MFG_L_7_2 | IFJ |
| | 190 | 18 | Frontal Lobe | Middle Frontal Gyrus | MFG_R_7_2 | IFJ |
| | 191 | 19 | Frontal Lobe | Middle Frontal Gyrus | MFG_L_7_3 | A46 |
| | 192 | 20 | Frontal Lobe | Middle Frontal Gyrus | MFG_R_7_3 | A46 |
| | 193 | 21 | Frontal Lobe | Middle Frontal Gyrus | MFG_L_7_4 | A9/46v |
| | 194 | 22 | Frontal Lobe | Middle Frontal Gyrus | MFG_R_7_4 | A9/46v |
| | 195 | 24 | Frontal Lobe | Middle Frontal Gyrus | MFG_R_7_5 | A8vl |
| | 196 | 28 | Frontal Lobe | Middle Frontal Gyrus | MFG_R_7_7 | A10l |
| | 197 | 29 | Frontal Lobe | Inferior Frontal Gyrus | IFG_L_6_1 | A44d |
| | 198 | 31 | Frontal Lobe | Inferior Frontal Gyrus | IFG_L_6_2 | IFS |
| | 199 | 32 | Frontal Lobe | Inferior Frontal Gyrus | IFG_R_6_2 | IFS |
| | 200 | 36 | Frontal Lobe | Inferior Frontal Gyrus | IFG_R_6_4 | A45r |
| | 201 | 46 | Frontal Lobe | Orbital Gyrus | OrG_R_6_3 | A11l |
| | 202 | 82 | Temporal Lobe | Middle Temporal Gyrus | MTG_R_4_1 | A21c |
| | 203 | 99 | Temporal Lobe | Inferior Temporal Gyrus | ITG_L_7_6 | A20cl |
| | 204 | 100 | Temporal Lobe | Inferior Temporal Gyrus | ITG_R_7_6 | A20cl |
| | 205 | 137 | Parietal Lobe | Inferior Parietal Lobule | IPL_L_6_2 | A39rd |
| | 206 | 138 | Parietal Lobe | Inferior Parietal Lobule | IPL_R_6_2 | A39rd |
| | 207 | 142 | Parietal Lobe | Inferior Parietal Lobule | IPL_R_6_4 | A40c |
| | 208 | 147 | Parietal Lobe | Precuneus | PCun_L_4_1 | A7m |
| | 209 | 148 | Parietal Lobe | Precuneus | PCun_R_4_1 | A7m |
| | 210 | 166 | Insular Lobe | Insular Gyrus | INS_R_6_2 | vIa |
| Default Mode (DM) | 211 | 3 | Frontal Lobe | Superior Frontal Gyrus | SFG_L_7_2 | A8dl |
| | 212 | 5 | Frontal Lobe | Superior Frontal Gyrus | SFG_L_7_3 | A9l |
| | 213 | 6 | Frontal Lobe | Superior Frontal Gyrus | SFG_R_7_3 | A9l |
| | 214 | 11 | Frontal Lobe | Superior Frontal Gyrus | SFG_L_7_6 | A9m |
| | 215 | 13 | Frontal Lobe | Superior Frontal Gyrus | SFG_L_7_7 | A10m |
| | 216 | 14 | Frontal Lobe | Superior Frontal Gyrus | SFG_R_7_7 | A10m |
| | 217 | 23 | Frontal Lobe | Middle Frontal Gyrus | MFG_L_7_5 | A8vl |
| | 218 | 33 | Frontal Lobe | Inferior Frontal Gyrus | IFG_L_6_3 | A45c |
| | 219 | 34 | Frontal Lobe | Inferior Frontal Gyrus | IFG_R_6_3 | A45c |
| | 220 | 35 | Frontal Lobe | Inferior Frontal Gyrus | IFG_L_6_4 | A45r |
| | 221 | 41 | Frontal Lobe | Orbital Gyrus | OrG_L_6_1 | A14m |
| | 222 | 42 | Frontal Lobe | Orbital Gyrus | OrG_R_6_1 | A14m |
| | 223 | 43 | Frontal Lobe | Orbital Gyrus | OrG_L_6_2 | A12/47o |
| | 224 | 44 | Frontal Lobe | Orbital Gyrus | OrG_R_6_2 | A12/47o |
| | 225 | 51 | Frontal Lobe | Orbital Gyrus | OrG_L_6_6 | A12/47l |
| | 226 | 52 | Frontal Lobe | Orbital Gyrus | OrG_R_6_6 | A12/47l |
| | 227 | 79 | Temporal Lobe | Superior Temporal Gyrus | STG_L_6_6 | A22r |
| | 228 | 80 | Temporal Lobe | Superior Temporal Gyrus | STG_R_6_6 | A22r |
| | 229 | 81 | Temporal Lobe | Middle Temporal Gyrus | MTG_L_4_1 | A21c |
| | 230 | 83 | Temporal Lobe | Middle Temporal Gyrus | MTG_L_4_2 | A21r |
| | 231 | 84 | Temporal Lobe | Middle Temporal Gyrus | MTG_R_4_2 | A21r |
| | 232 | 87 | Temporal Lobe | Middle Temporal Gyrus | MTG_L_4_4 | aSTS |
| | 233 | 88 | Temporal Lobe | Middle Temporal Gyrus | MTG_R_4_4 | aSTS |
| | 234 | 95 | Temporal Lobe | Inferior Temporal Gyrus | ITG_L_7_4 | A20il |
| | 235 | 121 | Temporal Lobe | Posterior Superior Temporal Sulcus | pSTS_L_2_1 | rpSTS |
| | 236 | 122 | Temporal Lobe | Posterior Superior Temporal Sulcus | pSTS_R_2_1 | rpSTS |

| | | | | | | |
|---|---|---|---|---|---|---|
| | 237 | 141 | Parietal Lobe | Inferior Parietal Lobule | IPL_L_6_4 | A40c |
| | 238 | 144 | Parietal Lobe | Inferior Parietal Lobule | IPL_R_6_5 | A39rv |
| | 239 | 153 | Parietal Lobe | Precuneus | PCun_L_4_4 | A31 |
| | 240 | 154 | Parietal Lobe | Precuneus | PCun_R_4_4 | A31 |
| | 241 | 175 | Limbic Lobe | Cingulate Gyrus | CG_L_7_1 | A23d |
| | 242 | 176 | Limbic Lobe | Cingulate Gyrus | CG_R_7_1 | A23d |
| | 243 | 179 | Limbic Lobe | Cingulate Gyrus | CG_L_7_3 | A32p |
| | 244 | 181 | Limbic Lobe | Cingulate Gyrus | CG_L_7_4 | A23v |
| | 245 | 187 | Limbic Lobe | Cingulate Gyrus | CG_L_7_7 | A32sg |
| | 246 | 188 | Limbic Lobe | Cingulate Gyrus | CG_R_7_7 | A32sg |
